# Parasitism and locomotory capacity calibrate the mitogenomic evolutionary rates in Bilateria

**DOI:** 10.1101/2023.02.28.530434

**Authors:** Ivan Jakovlić, Hong Zou, Tong Ye, Hong Zhang, Xiang Liu, Chuan-Yu Xiang, Gui-Tang Wang, Dong Zhang

## Abstract

The evidence that parasitic lineages exhibit elevated evolutionary rates is limited to Arthropoda and inconsistent. Similarly, the evidence that mitogenomic evolution is faster in species with low locomotory capacity (LC) is limited to a handful of animal lineages. We hypothesised that these two variables are associated and that LC is a major underlying factor driving the elevated evolutionary rates in parasites. We tested this hypothesis by studying mitogenomic evolutionary patterns in 10,911 bilaterian species classified according to their locomotory capacity and parasitic/free-living life history (LH). Evolutionary rates were significantly elevated in endoparasites, ectoparasites with reduced LC, and free-living lineages with reduced LC, but not in ectoparasites and parasitoids with high LC. Nematoda and Arachnida were the only lineages where parasitism was not associated with faster evolution. We propose that LC may also explain these two major outliers. Overall, the LH categorisation explained 35-37%, LC categorisation 26-28%, and together they explained 41-44% of the variance in branch lengths across the Bilateria. Our findings suggest that these two variables play a major role in calibrating the molecular clock in bilaterian animals.

## Introduction

Rates of mitogenomic (mitochondrial genome) molecular evolution are remarkably variable among animal lineages. While studies identified numerous factors that may be associated with comparatively elevated mitogenomic evolutionary rates, their effects appear to be small, inconsistent, lineage-specific, and mitogenomic evolution thus very difficult to predict ^1–5^. Several studies found evidence that parasitism might be associated with elevated mitochondrial sequence evolution rates ^6–10^. However, all of these studies were conducted on different Arthropoda lineages. As regards non-arthropod animals, there is merely anecdotal evidence. Parasitic and/or commensal Clitellata (Annelida) appear to exhibit elevated evolutionary rates ^11^, but this dataset was small and the finding was not tested statistically. In a meta-analysis of all bilaterian mitogenomes available in 2012, Bernt et al. observed that parasitism exhibited a somewhat patchy relationship with increased branch lengths ^12^, but did not test this statistically either. They found exceptionally long branches in Nematoda, Platyhelminthes, Tunicata, some Mollusca, and some Arthropoda (Acari and Copepoda). Most of these taxa comprise both free-living and parasitic lineages, but some comprise only parasitic and some only free-living species. Nematodes, which exhibit a complex mix of parasitic and free-living life histories ^13^, are known to possess rapidly evolving mitogenomes ^14,15^, but Bernt et al. concluded that there appears to be no clear correlation between the branch length and parasitic lifestyle in this clade ^12^. Over the last 6-7 years, we sequenced over 30 mitogenomes of parasitic animals, largely Platyhelminthes and Nematoda, but also some Arthropoda (Supplementary file S1: Table S1). We found some indirect evidence that parasites may be disproportionately likely to have rapidly evolving and architecturally destabilised mitogenomes ^16–18^, but also did not test this statistically.

Furthermore, even in arthropods, the association between parasitism and evolutionary rate is inconsistent: in insects, parasitism was associated with an increased evolutionary rate in Hymenoptera, but not in Diptera ^19^. Therefore, the association between elevated mitochondrial sequence evolution rates and parasitism is inconsistent (lineage-specific), and it remains statistically untested for taxa other than certain arthropod lineages to our knowledge. Further studies are needed to test the universality of this phenomenon in animals.

The underlying cause for this putative correlation between elevated mitogenomic evolutionary rates and parasitism also remains unresolved. Previously proposed candidates include directional selection driven by the genetic arms race between hosts and parasites ^6,20^, the compensation-draft feedback ^20^, or the reduction of purifying selection pressures in parasitic species, which may be driven by the effective population size (*N*_*e*_) reductions ^21^ driven by high speciation rates in parasites and frequent founder events during transmissions to new host individuals ^6,19,22^. Following the evidence that purifying selection pressure is correlated with the section for locomotory capacity in crustaceans ^10^, we hypothesise that relaxation of purifying selection pressures in (some, but not all) parasitic lineages may be further driven by a strongly reduced locomotory capacity. The relaxation may be additionally driven by the overall reduction of various biological processes in some parasitic lineages ^23,24^. The underlying logic is that mitochondrial genomes play a central role in the production of energy, so the strength of purifying selection acting on mitochondrial genes should be positively correlated with the energy expenditure, which in turn should produce a positive correlation with the strength of selection for locomotory capacity ^10^. In endoparasites and some ectoparasites, an almost complete absence of locomotory capacity, putatively in combination with high metabolic dependence on the host, may allow for a certain level of degradation of the mitogenomic energy production efficiency. The existence of a positive correlation between the metabolic rate and purifying selection pressures was observed in salamanders ^25^ and crustaceans ^10^, whereas the association between the locomotory capacity and the strength of purifying selection on the mitochondrial genome was confirmed in molluscs ^26^ and crustaceans ^10^, as well as selected insect ^27^, fish ^28,29^, bird, and mammal ^30^ lineages. However, this hypothesis was never comprehensively tested on a broader dataset, and parasitism and locomotory capacity were never associated by previous studies to our knowledge.

We hypothesise that locomotory capacity may be the crucial factor that explains why the association between parasitism and elevated mitogenomic evolutionary rates appears to be patchy. While the role of the *N*_*e*_ in this inconsistency has been recognised previously ^19^, the locomotory capacity appears to have been overlooked by previous studies. Herein we adopted the definition of parasitism as a consumer interaction in which the consumer feeds on a single individual (the host) during at least one life-history stage, where both parasites and hosts belong to the Animalia, which includes strategies employed by parasitoids, parasitic castrators, macroparasites and pathogens, but excludes micropredators, brood ‘parasites’, kleptoparasites, symbiotic egg predators, inquilines, non-feeding symbionts, and plant-parasites ^31,32^. Selection for locomotory capacity is not uniform across different parasitic strategies: whereas endoparasites generally possess merely a rudimentary locomotory capacity, certain ectoparasitic (such as fleas) and most parasitoid lineages are evolving under a comparatively high selection for locomotory capacity. As a result, mitogenomes of parasitoid organisms, which commonly only have a parasitic larval stage and free-living adult stage, must evolve under stringent purifying selection pressures, because degradation of the mitogenomic energy production efficiency would affect the fitness of free-living adults. Additionally, due to the existence of free-living adults, there is no reason to expect a reduced *N*_*e*_ in comparison to fully free-living species in parasitoids. Contrary to this, most endoparasitic species, even when their life history includes multiple hosts, do not undergo selection for locomotory capacity during any of their life stages. Along with the high metabolic dependence on the host, this may allow strongly reduced metabolic rates, and consequently relaxation of purifying selection pressures on mitochondrial genes. Purifying selection may be additionally weakened by a reduced *N*_*e*_, which is expected to be propelled by the high speciation rate, common asexual reproduction, and population fragmentation (reproducing adults are confined to the host) ^33^.

Following this logic, we would expect endoparasites to exhibit the highest evolutionary rates, followed by ectoparasitic lineages exhibiting low locomotory capacity, but we would not expect parasitoids to differ significantly from free-living lineages. As several previous studies were referring to parasitoids when identifying elevated mitogenomic evolutionary rates in parasites ^6,19,20^, this would explain the inconsistent results ^19^, and suggest that elevated evolutionary rates may have been mistakenly attributed to ‘parasitism’, when in fact they were driven by other variables.

To test these hypotheses, in this study we used all Bilateria mitogenomes available in the non-redundant RefSeq database ^34^ (last accessed on 10^th^ March 2022). We divided parasitic life histories into endoparasites (comprising mesoparasites) (EndoP), ectoparasites (EctoP), parasitoids, and species that alternate between parasitic and free-living (F) generations (EndoP/F). To further confirm our predictions, we separately classified micropredators (MP), which are treated as parasites in some classifications ^35^. They resemble many ectoparasites in their haematophagous feeding habits, but they only visit the host for feeding, and otherwise they are free-living (this category comprises animals such as mosquitoes and some bats). We hypothesised that their evolutionary rates should not differ from other free-living organisms. We also roughly divided the entire dataset according to the locomotory capacity into three categories: low, intermediate, and high. We hypothesised that: 1. endoparasitic lineages would exhibit the fastest evolutionary rates due to almost complete loss of locomotory capacity, metabolic dependence on the host, and physical confinement to the host; 2. ectoparasites would produce a lineage-specific, noisy signal, because in terms of locomotory capacity, metabolic dependence on the host, and physical confinement, certain ectoparasitic lineages (such as flea) are similar to many free-living and parasitoid lineages, whereas others (such as monogenean flatworms) are relatively similar to endoparasites; 3. evolutionary rates should not differ between the free-living, parasitoid and micropredator lineages; 4. lineages with low locomotory capacity should exhibit increased evolutionary rates regardless of their life history. We tested these hypotheses and compared the explanatory powers that the two variables have on mitogenomic evolution in Bilateria using the complete bilaterian dataset as well as a large number of subsets designed to control for different confounding factors.

## Results

### Pairwise comparisons at different taxonomic levels

The final dataset comprised 10,911 species classified into 24 phyla, but a majority were Chordata (6,228) and Arthropoda (3,504) (Supplementary file S2: Worksheet A). A vast majority were classified as free-living, but the numbers of parasitoid, micropredatory, ectoparasitic, and endoparasitic species were sufficient for statistical analyses (64 to 275). Only five EndoP/F species (Strongyloididae nematodes alternating between free-living and parasitic generations) had weak statistical power (Supplementary file S1: Table S2). Accounting for some uncertainty in evolutionary scenarios, we counted around 6 independent origins of endoparasitism and 12 origins of ectoparasitism in our dataset. Average branch lengths were the highest in Platyhelminthes (3.7, 92% of parasitic species), followed by Orthonectida (3.0, only one species, endoparasite *Intoshia linei*), Acanthocephala (2.9; 100% endoparasitic), and Nematoda (2.5, 81% parasitic) (Table 1, Figure 1). The only phylum containing parasitic species that did not exhibit exceptionally long braches was Arthropoda, but the ratio of parasites was low (2.9%). Endoparasites exhibited by far the longest average branches in the dataset (3.11), almost twice (1.82×) as long as the free-living species. They were followed by the EndoP/F group (2.38), ectoparasites (2.01), parasitoids (1.43), and finally the shortest and almost identical branches in micropredators (1.26) and free-living species (1.25) (Figure 1). Differences were non-significant among the latter three groups using Tukey HSD tests (Figure 2; Supplementary file S1: Table S1, exact p-values in Supplementary file S2: Worksheet B). Endoparasites and ectoparasites had much higher SD (standard deviation) values (0.7-0.9) than other groups (>0.3), but the effect sizes were large for comparisons of both groups to the free-living (Supplementary file S1: Table S2 and Figure S1). Using the phylogenetic generalised least squares (PGLS) ANOVA test, we found significant differences in all comparisons (pairwise and overall) and all datasets, regardless of how small the differences were (Supplementary file S1: Figure S2), so we suspect that it overestimates the statistical significance of differences. We therefore refer to Tukey HSD tests when discussing the significance of differences. As multiple categories exhibited outliers (Figure 2), we also corroborated that these relationships are stable when outliers are removed: the overall patterns were upheld, and most of the main conclusions were further strengthened (Supplementary file S1: Figure S3). We further assessed whether these patterns are consistent at lower phylogenetic levels. They were closely mirrored in Ecdysozoa, Lophotrochozoa, Ecdysozoa+Lophotrochozoa, Platyhelminthes and Arthropoda clades, with the exception of some differences in p-values, and endoparasitic and ectoparasitic lineages exhibiting identical average branch lengths in Lophotrochozoa (Supplementary file S1: Text S1). A major exception was Nematoda, where branch lengths of endoparasitic lineages were only nonsignificantly longer than in the free-living. Notably, all species in this phylum were assigned to the low LC group. At lower taxonomic levels (class and order), results were somewhat less consistent than at higher levels, but analyses were weakened by low numbers of samples for many categories. Among datasets with sufficient statistical power, Insecta (class) and Diptera (order) were in full agreement with working hypotheses, whereas in the class Arachnida, ectoparasites had shorter branches than the free-living (Supplementary file S1: Text S2 and S3, and Figures S4 to S9).

**Table 1.**
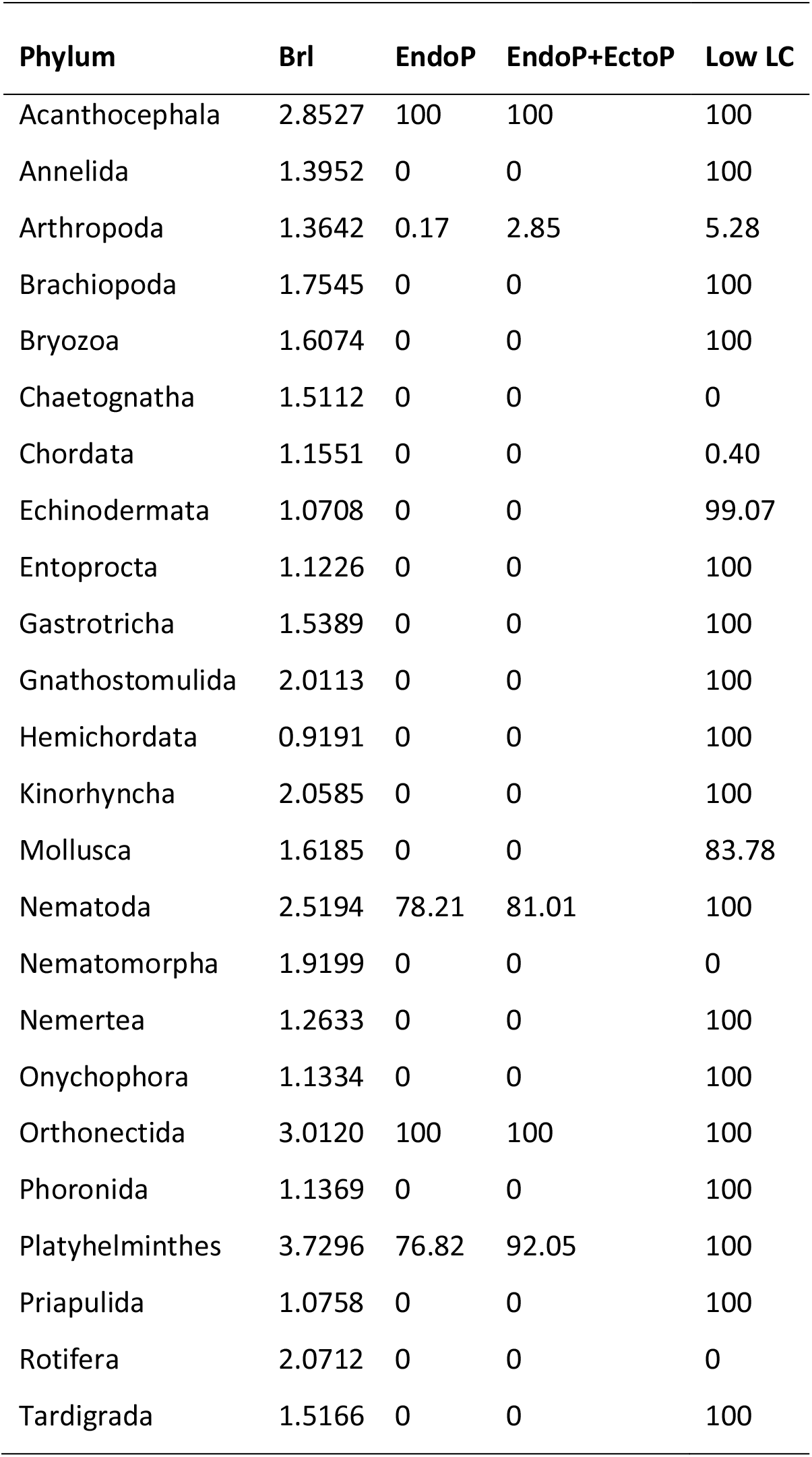
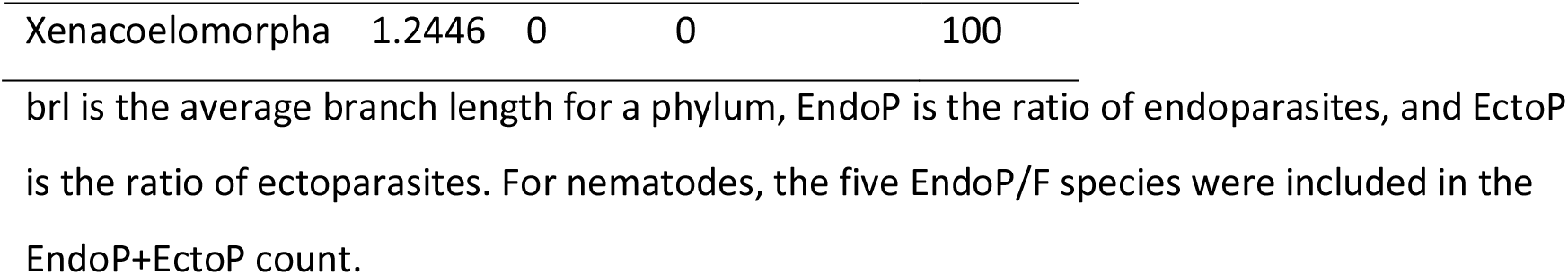
The ratios of parasitic and low locomotory category (Low LC) species in each phylum represented in the dataset.

| Phylum | Brl | EndoP | EndoP+EctoP | Low LC |
| --- | --- | --- | --- | --- |
| Acanthocephala | 2.8527 | 100 | 100 | 100 |
| Annelida | 1.3952 | 0 | 0 | 100 |
| Arthropoda | 1.3642 | 0.17 | 2.85 | 5.28 |
| Brachiopoda | 1.7545 | 0 | 0 | 100 |
| Bryozoa | 1.6074 | 0 | 0 | 100 |
| Chaetognatha | 1.5112 | 0 | 0 | 0 |
| Chordata | 1.1551 | 0 | 0 | 0.40 |
| Echinodermata | 1.0708 | 0 | 0 | 99.07 |
| Entoprocta | 1.1226 | 0 | 0 | 100 |
| Gastrotricha | 1.5389 | 0 | 0 | 100 |
| Gnathostomulida | 2.0113 | 0 | 0 | 100 |
| Hemichordata | 0.9191 | 0 | 0 | 100 |
| Kinorhyncha | 2.0585 | 0 | 0 | 100 |
| Mollusca | 1.6185 | 0 | 0 | 83.78 |
| Nematoda | 2.5194 | 78.21 | 81.01 | 100 |
| Nematomorpha | 1.9199 | 0 | 0 | 0 |
| Nemertea | 1.2633 | 0 | 0 | 100 |
| Onychophora | 1.1334 | 0 | 0 | 100 |
| Orthonectida | 3.0120 | 100 | 100 | 100 |
| Phoronida | 1.1369 | 0 | 0 | 100 |
| Platyhelminthes | 3.7296 | 76.82 | 92.05 | 100 |
| Priapulida | 1.0758 | 0 | 0 | 100 |
| Rotifera | 2.0712 | 0 | 0 | 0 |
| Tardigrada | 1.5166 | 0 | 0 | 100 |
| Xenacoelomorpha | 1.2446 | 0 | 0 | 100 |

brl is the average branch length for a phylum, EndoP is the ratio of endoparasites, and EctoP is the ratio of ectoparasites. For nematodes, the five EndoP/F species were included in the EndoP+EctoP count.

**Figure 1.**
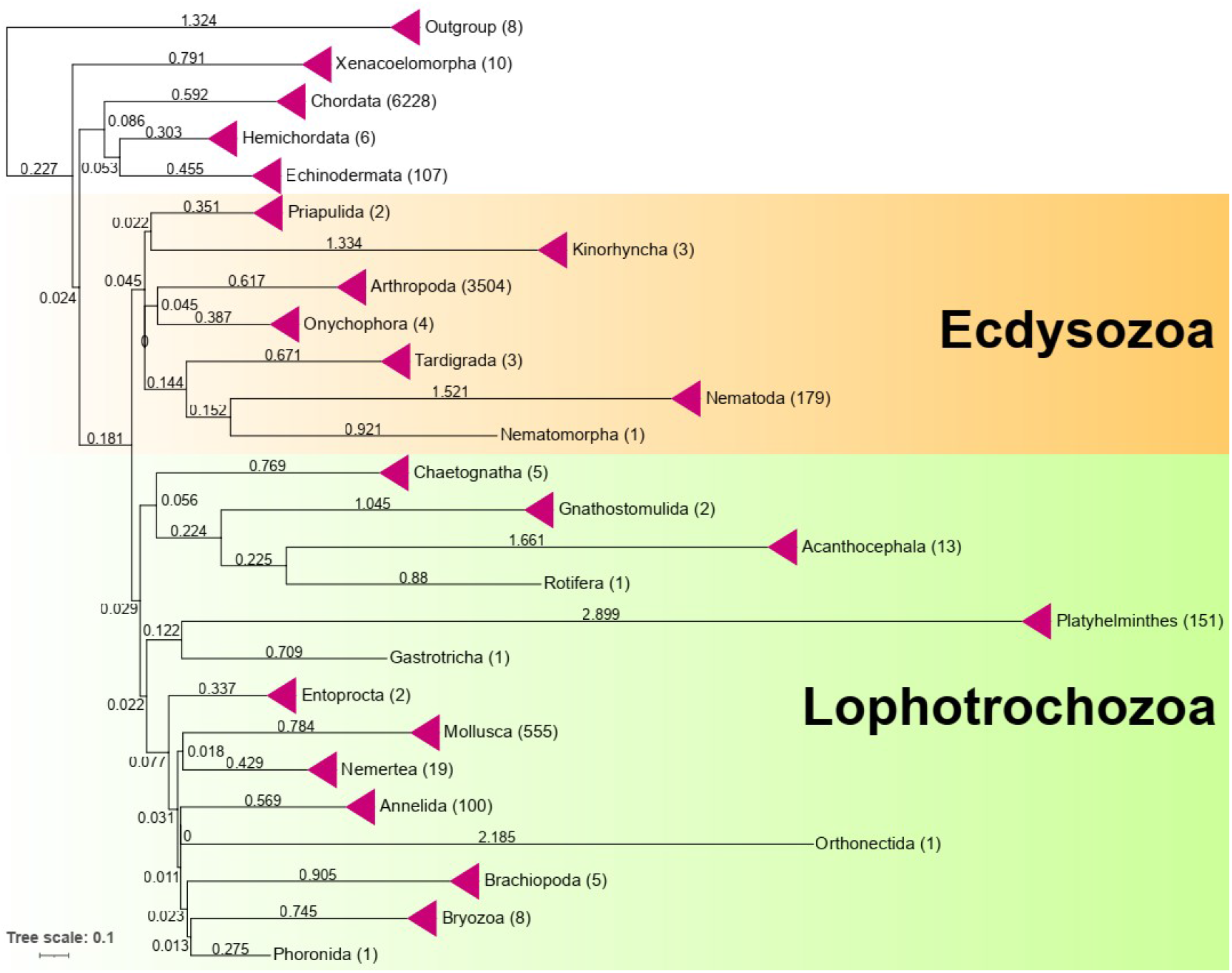
Phylogeny of Bilateria topology-constrained at the phylum level, inferred using amino acid sequences of 12 mitochondrial protein-coding genes. Branch lengths are shown in the tree. All phyla were collapsed to a single branch. The number in the parentheses next to the leaf name (phylum) denotes the number of species in the clade.

**Figure 2.**
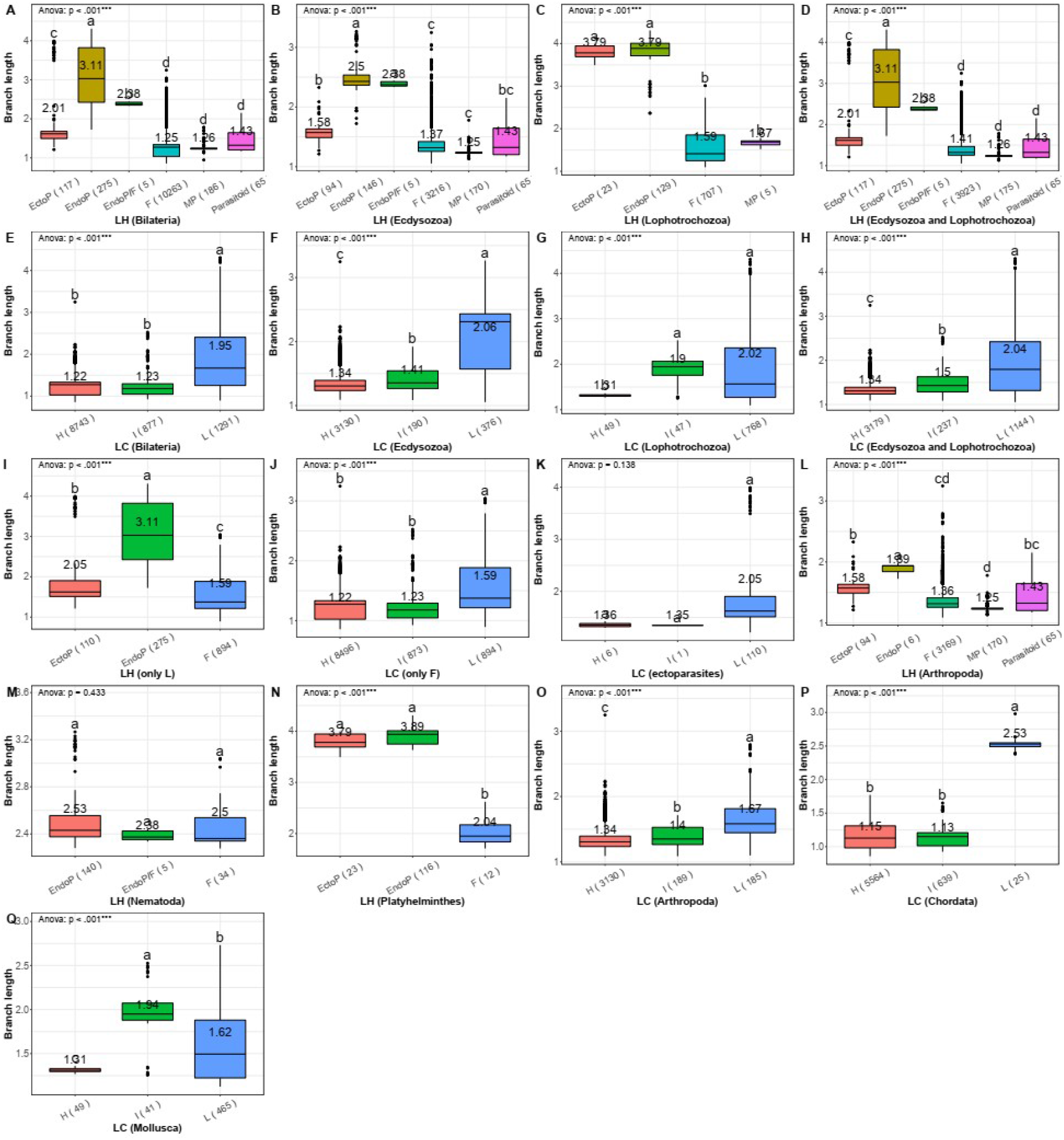
Branch length comparison between different life history and locomotory capacity categories. In the life-history categorisation (LH), F is free-living, EndoP is endoparasitic, EctoP is ectoparasitic, MP is micropredatory, and EndoP/F refers to interchanging free-living and parasitic generations. In the locomotory capacity categorisation (LC), H is high, I is intermediate, and L is low. The dataset on which the analysis was conducted is indicated beneath each chart. Boxplots show the minimum, first quartile, median, third quartile, and maximum, plus outliers. Average branch length values are shown in the graphs. PGLS ANOVA results are shown in the upper left corner. Different letters above the boxplots indicate statistically significant differences (p<0.05; Tukey HSD). The number of species included in the analysis is shown next to the category name (x-axis).

As regards the locomotory capacity (LC) classification, the vast majority of Bilateria were classified as ‘High’, but both remaining categories comprised over 800 species, ensuring good statistical power. In agreement with our working hypothesis, branch lengths were the lowest in the High and Intermediate LC categories (1.22 and 1.23 respectively), and much higher (significantly) in the Low LC category (1.95) (Figure 2, Supplementary file S1: Table S1). The Low category had a much larger SD value (0.9) than the other two groups (≈0.2), and the effect sizes were medium (Low and Intermediate groups vs. the High group; Supplementary file S1: Table S1 and Figure S1). Branches were shorter in the high LC category than in the low LC category in all lower-level taxa where both categories were included, but the intermediate category produced a somewhat noisy pattern (additional details in Supplementary file S1: Text S1 and Figures S4 to S9). Within the free-living dataset (>10,000 species), among the 200 species with the highest branch lengths, 186 were classified in the Low LC category, and only 5 in the High LC category. In addition, we also confirmed that single-gene topologies produce congruent results (Supplementary file S1: Text S4).

As parasitism and locomotory capacity are partially overlapping variables, we attempted to discern their impacts using subsets of data. To reduce the effect of locomotory capacity variability, we focused only on the Low LC category. The pattern produced by the overall bilaterian dataset was upheld, but the average branch lengths were increased compared to the overall dataset from 2.01 to 2.05 in ectoparasites and from 1.25 to 1.59 in free-living species (Figures 2A and 2I). As all endoparasites were classified as the Low LC category, their branch length was unaffected (3.11). To remove the effect of parasitism, we conducted analyses using only the free-living species. Again, the pattern was upheld, with branch lengths unchanged in the High (1.22) and Intermediate (1.23) categories, but much shorter in the Low LC category (1.59 vs. 1.95) (Figures 2E and 2J). We further focused on the ectoparasitic dataset and compared lineages according to the LC classification: species in the High (1.35) and Intermediate (1.36) categories (all Arthropoda and Insecta) had much shorter branches than those in the Low category (2.05) (Figure 2K). Moreover, their branch lengths were almost identical to those in free-living species with High LC in Arthropoda and Insecta (both ≈1.35; Figure 2 and Supplementary file S1: Figures S7 and S11). The statistical power of these analyses was weakened by a low number of species in the Low and Intermediate ectoparasite categories.

### Multiple regression and LRT analyses

The *lmekin* (multilevel phylogenetic regression) analysis of the bilaterian dataset divided into six LH categories and three LC categories found that these two categorisations together explained 44% of the branch length variance in the dataset. Individually, the two variables explained 37% (LH) and 28% (LC) of the variance (Table 2). We tested whether dividing the dataset differently along the life history classification lines (e.g. EndoP+EctoP vs. all other categories) affects the results, but R^2^ values were largely unaffected (Supplementary file S1: Table S5). Removing the phylogenetic signal by running a lmekin analysis without importing the phylogenetic matrix produced a negligible difference in R^2^ values. We also confirmed these results using *brms*: results were very similar, but R^2^ values were on average 2-4% lower than those inferred by *lmekin* (Supplementary file S1: Table S6). To reduce the collinearity between parasitism and the low LC category, we tested the explanatory power of the LC categorisation using only the free-living dataset: LC explained ≈ 11% of the variance in branch lengths (Table 2). Similarly, we conducted an analysis using only the low LC category to test the impact of LH categorisation (EndoP, EctoP, MP, F): it explained over 50% of the branch length variance (Table 2). We designed likelihood-ratio tests (LRT) to assess the explanatory power of life history and locomotory capacity categorisation in relation to the branch length. In all cases, these categorisations had a better model fit (lower AIC value) and greater explanatory power (higher R^2^ value) than the null model (includes only random effects and correction for phylogeny), and implementation of both variables had a better model fit and greater explanatory power than either of the variables alone (Table 3).

**Table 2.**
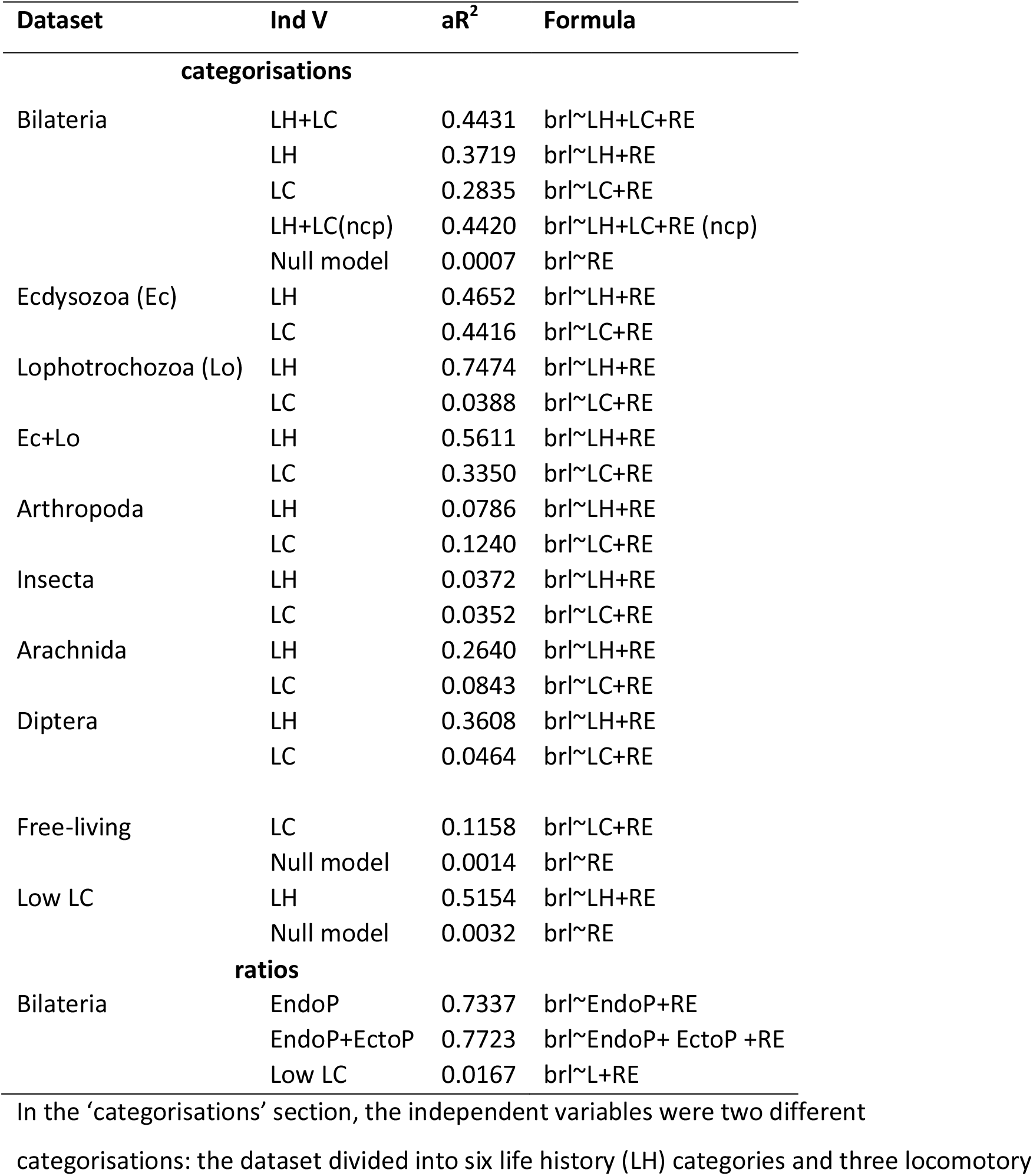
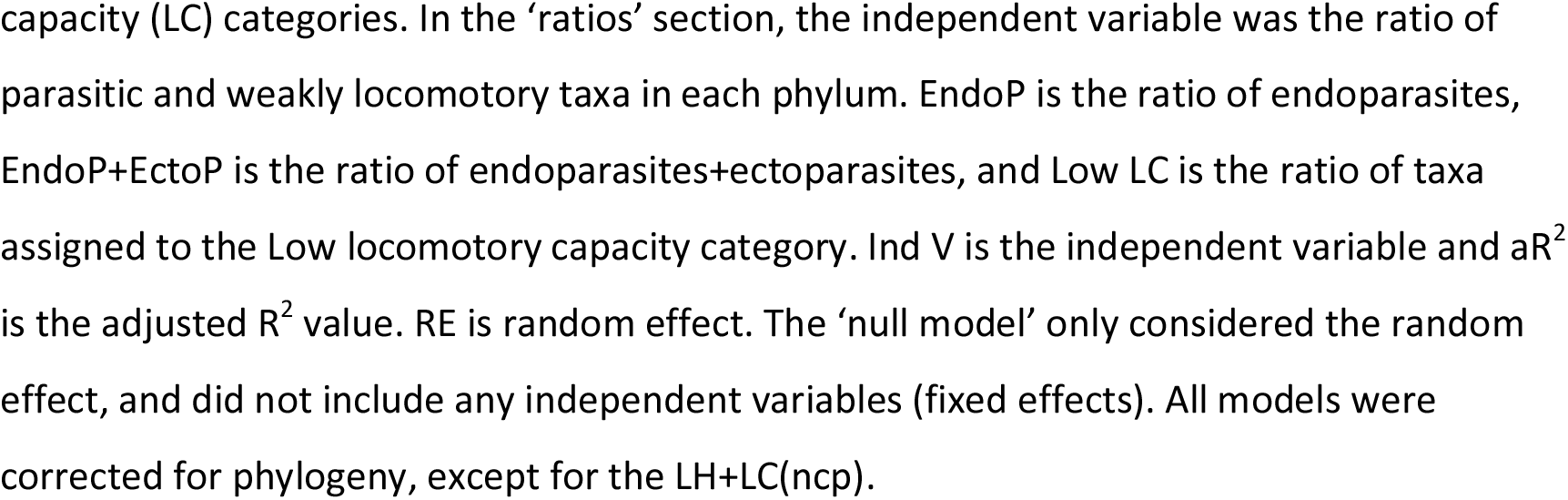
Phylogeny-corrected *lmekin* multilevel regression analyses with branch length as the dependent variable.

**Table 3.**
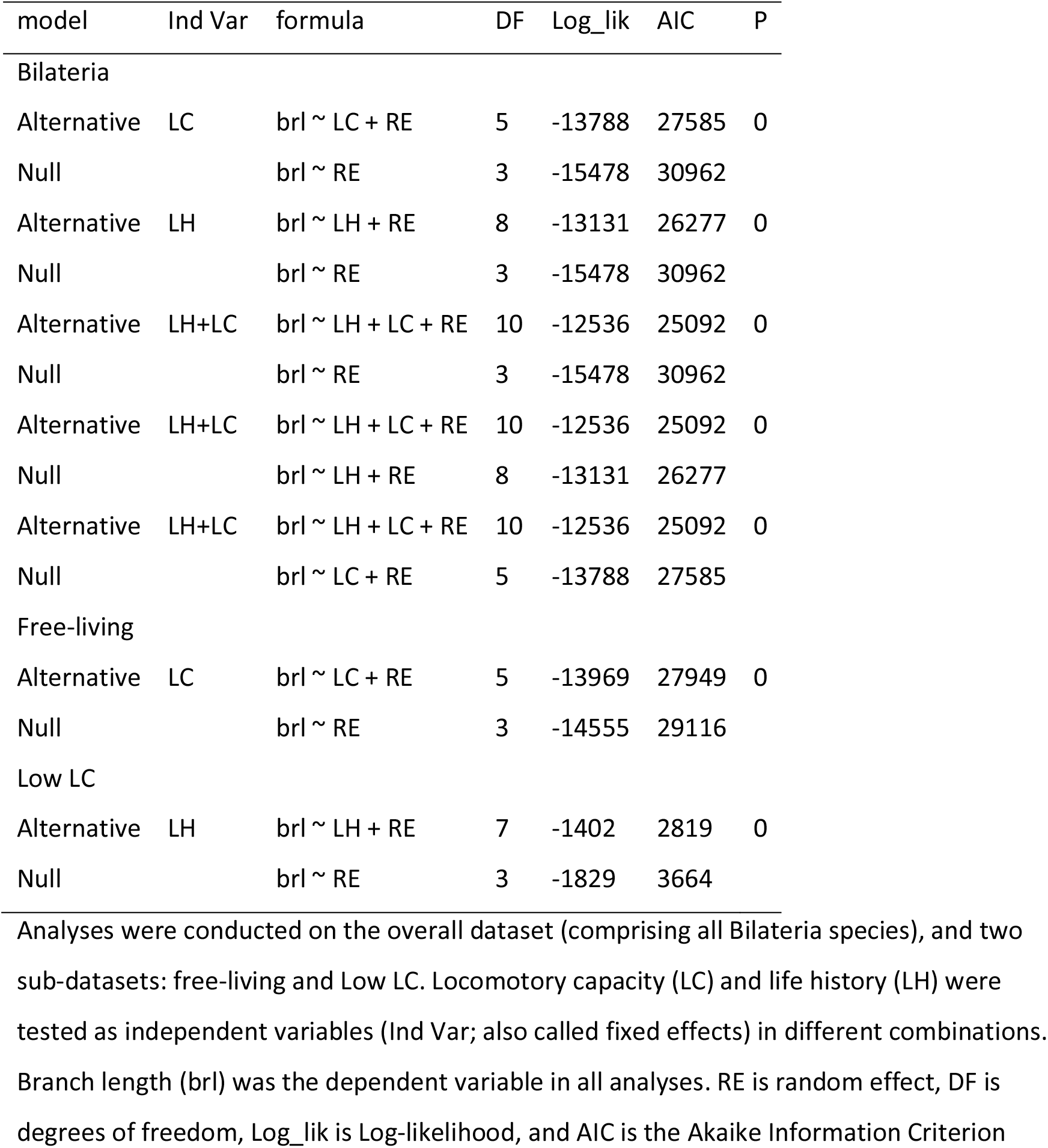
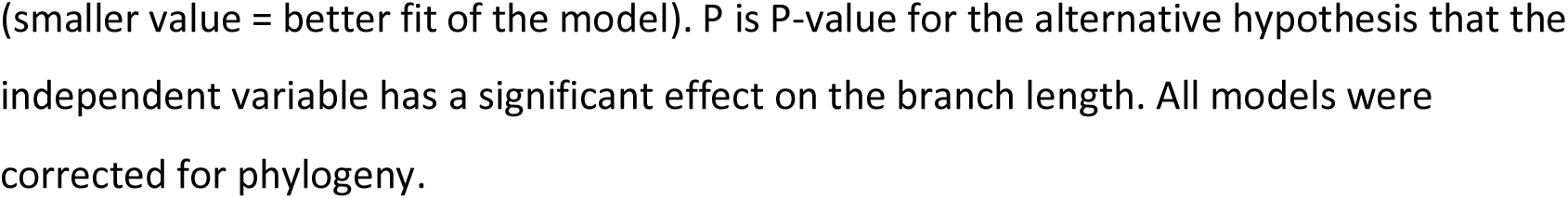
Likelihood Ratio Test (LRT) designed to assess the impact of life history and locomotory capacity on the branch length.

To test for lineage-specific effects, we conducted *lmekin* analyses on several taxonomic lineages that contained a mix of LH and LC categories. In the Ecdysozoa, life history and locomotory capacity categorisations explained almost half of the variance each (≈45%) (Table 2). The subset Lophotrochozoa exhibited a unique pattern: life history explained a major proportion of the variance (75%), whereas locomotory capacity explained a small amount of it (4%). The Ecdysozoa+Lophotrochozoa dataset produced an intermediate result. At the phylum level, locomotory capacity explained a greater (12%) proportion of the variance than life history (8%) in Arthropoda. At lower levels, both variables explained a relatively small amount of variance (≈4%) in Insecta; in Arachnida and Diptera, life history explained 26 to 36% of the total variance, whereas locomotory capacity explained less than 10% (5-8%).

The ratio of endoparasites in a phylum explained 73.3% of the branch length variance. The addition of ectoparasites (EndoP and EctoP) slightly increased the explanatory power (77.2%). The ratio of species assigned to the low LC group had a small explanatory power (1.8%) (Tables 1 and 2).

### Selection pressure analyses

We further tested the hypothesis that parasitic and low LC lineages exhibit signals of relaxed purifying selection pressures. These analyses had to be conducted on strongly reduced datasets due to computational limitations and the existence of multiple genetic codes within the Bilateria. The purifying selection pressures were relaxed in endoparasites and parasitoids (but not in ectoparasites) vs. the free-living, and in Low vs. High LC groups. We also tested for directional selection: there were significant signals in all groups in most cases. Overall, these results were rather noisy (details in Supplementary file S1: Text S5 and Tables S7 and S8).

## Discussion

We found consistent support for our working hypotheses at higher taxonomic levels. Although patterns were less consistent at lower phylogenetic levels, in both major outliers (Nematoda and Arachnida), the discrepancy can be at least partially explained by the locomotory capacity variable. In Nematoda, branch lengths of endoparasitic lineages were only nonsignificantly higher than those of the free-living, but the entire phylum exhibits a limited locomotory capacity, with a relatively small variation between the parasitic and free-living lineages. In Arachnida, ectoparasitic lineages (ticks) had shorter branches than the free-living ones (spiders, scorpions, mites etc.), but this was a product of exceptionally long branches in free-living mites, which possess merely a rudimentary locomotory capacity, comparable to those of parasitic lineages (Supplementary file S1: Text S2). Combined, these findings support the hypothesis that the reduction of locomotory capacity is among the key variables behind the elevated evolutionary rates in parasitic lineages. However, as the proportion of variance explained by the LC classification was on average smaller, and in some cases much smaller, than the proportion of variance explained by the life history classification, this indicates that locomotory capacity is not the only explanatory variable for elevated evolutionary rates in parasitic lineages. For example, locomotory capacity is also limited (albeit less so than in parasites) in free-living Platyhelminthes, along with a range of other convergent evolutionary trends between the two groups ^23^, but the evolutionary rates are highly elevated in parasitic compared to the free-living lineages.

Our findings indicate that answering the question ‘why?’ is important when attempting to answer the question: ‘does parasitism cause elevated evolutionary rates?’. This allowed us to predict and prove that only some parasitic life histories should exhibit significantly elevated evolutionary rates, which may help us explain inconsistent results in previous studies. Specifically, previous studies treated parasitoid insects as parasites, but herein we argued that because adults are free-living, they are evolving under selection for high locomotory capacity. In addition, adults are not physically confined to the body of the host, and they cannot be metabolically dependent on the host throughout their entire life, so they evolve under stronger purifying selection pressures than most macroparasites. However, parasitoids did exhibit nonsignificantly longer branches than free-living organisms, and selection analyses found signals of relaxed purifying selection pressures and elevated rates of directional selection in parasitoid organisms (compared to free-living). Assuming that these findings are not artefactual, they may explain the elevated evolutionary rates in some mitochondrial genes in hymenopteran parasitoid wasps ^6,19^. The factors behind these observations remain conjectural, but their impacts are too small to produce consistently significant results.

Importantly, *lmekin* and *brms* analyses indicate that life history categorisation (parasitism) explained 35-37%, LC categorisation 26-28%, and together they explained 41-44% of the variance in branch lengths across the Bilateria. This is much more than the generation time, which explained about 13% of the variability in mitogenomic evolutionary rates in invertebrates ^36^, and more than a broad range of life history and ecological variables in mammals ^2^. As parasitism and locomotory capacity overlap to an extent (most but not all parasitic lineages exhibit low LC), to remove the effect of parasitism, we conducted additional analyses using only free-living species. This reduced the proportion of the variability in branch length across the Bilateria explained by this variable (from 26-28% to 10-12%). The explanatory power of this variable was even lower in some sub-datasets, but it is unclear whether this is attributable to insufficient sampling, imprecise classification, or genuinely low explanatory power of this variable in certain lineages. In other aspects, this variable seemed to have remarkably good predictive power. For example, ectoparasites with high LC had the same branch lengths as free-living species with high LC, and extreme branch lengths within the free-living dataset were almost exclusively found in species with low locomotory capacity. We also attempted to minimise the effect of locomotory capacity variability by focusing on the Low LC category, which increased the already high proportion of variance explained by the LH categorisation from 37% to 51%. While these findings indicate that locomotory capacity is a variable driving the heterogeneity in evolutionary rates in bilaterian animals, as well as elevated rates in parasitic lineages, we failed to prove the hypothesis that it is the most important variable. Other variables also play a role, and in some cases even outweigh the impact of locomotory capacity on mitogenomic evolution. These comprise metabolic dependence on the host, directional selection, effective population size, generation time, replication and repair machinery, etc. (further discussion in Supplementary file S1: Text S6).

Due to its much broader dataset scope, the resolution of our study is much higher than in previous attempts to tackle this question, but there are several limiting factors. A potential source of noise in our analyses lies in the fact that many nominally parasitic species have free-living stages ^32^, and that boundaries between different parasitic life history categories, and even free-living and parasitic ^23^, are often blurry. For example, all plant parasites were included in the free-living category, although some may be similar to those classified as parasitic in terms of reduced locomotory capacity and confinement to a single host. There was even more noise in the locomotory capacity categorisation, as there are no clear lines according to which species can be categorised in this aspect, so our categorisation of locomotory capacity was insufficiently precise to capture the full range of differences in locomotory capacity between bilaterian species. There is also some overlap between the three LC categories, but the goal was to minimise the overlap between the High and Low LC categories. This was achieved by introducing the Intermediate category, wherein we put all species expected to exhibit relaxed locomotory capacity selection pressures, but insufficiently so to be included in the Low group. In this way, both categories probably exhibit some overlap with the Intermediate group, but the overlap between L and H should be minimal. Due to the weak resolution of this categorisation, it is likely that the explanatory power of this variable was underestimated in our study. As our dataset did not conform to the assumption of independence of data (phylogenetic relatedness), for pairwise analyses, we conducted both ordinary (Tukey HSD) and phylogeny-corrected statistical tests (PGLS ANOVA). We found evidence that PGLS ANOVA may have overestimated the statistical significance of differences. This has been observed before, and it was proposed that in such cases traditional non-phylogenetically controlled approaches might be statistically more appropriate ^37^, so we largely relied on the standard Tukey HSD test. In addition, multilevel regression analyses were barely affected by accounting for the phylogenetic relatedness of data. Additional confounding variables are discussed in Supplementary file S1: Text S7.

## Summary and conclusions

We showed herein that previous evidence that parasitic lineages exhibit elevated evolutionary rates in comparison to free-living lineages is largely limited to selected Arthropoda lineages, that results are often inconsistent, and we argued that there is no explanation for why some nominally parasitic life history strategies, such as parasitoids, would exhibit elevated evolutionary rates, thus leaving the question unresolved. To attempt to get a more or less definitive answer to this problem, we analysed almost 11,000 available bilaterian mitogenomes, and provided the first statistical evidence that elevated mitogenomic evolutionary rates in endoparasitic and most ectoparasitic lineages are an almost universal phenomenon in Bilateria. Across the Bilateria, endoparasitic lineages consistently (nematodes aside) exhibit much higher evolutionary rates than free-living lineages. Ectoparasites also exhibited strongly elevated evolutionary rates, but less pronounced than endoparasites. In partial agreement with our predictions, parasitoids and micropredators exhibited non-significantly elevated evolutionary rates. We also found support for the hypothesis that evolutionary rates are higher in bilaterians with a strongly reduced locomotory capacity. The explanatory power of locomotory capacity varied between lineages and analyses. It remains unclear whether this was due to the imprecision of our classification, or the fact that other variables outweigh its impacts. The selection for locomotory capacity does appear to be a suitable explanation for the major outliers with respect to our hypotheses (Nematoda and Arachnida), and parasitic life histories with comparatively high locomotory capacity, a subset of ectoparasites (fleas) and parasitoids, had much lower evolutionary rates than parasites with low locomotory capacity. However, endoparasites, followed by some ectoparasites (Monogenea) may also exhibit the highest level of metabolic dependence on the host and strongly reduced *N*_*e*_ values, so we must account for the possibility that other variables overlap with the locomotory capacity. While our analyses support previous conclusions that there is no single factor with the power to predict the patterns of mitogenomic evolution in all animal lineages ^1,38,39^, life history categorisation (parasitism) explained 35-37% of the variance, locomotory capacity categorisation explained 26-28%, and together they explained 41-44% of the total variability in evolutionary rates of mitochondrial PCGs in Bilateria, which indicates that we managed to identify the major explanatory variables for mitogenomic evolution in bilaterian animals.

## Materials and Methods

### Dataset

When we accessed the data (10^th^ March 2022), there were 11,284 animal and 11,017 bilaterian mitogenomes available in the RefSeq database. What is commonly recognised as the ‘standard’ metazoan (animal) mitogenome is a circular molecule ≈15 Kbp in size that contains 37 genes: 13 protein-coding genes (PCGs), 2 rRNA genes and 22 tRNA genes, but there are major deviations from this architecture in some lineages. While some deviations from this canon have been observed in isolated bilaterian lineages, almost all of the major discrepancies in the protein-coding gene content map to the non-bilaterian metazoans ^40^. As this would complicate some comparative analyses, and as non-bilaterian mitogenomes represent only about 2% of the total available Animalia dataset, in this study we focused only on Bilateria. PhyloSuite ^41^ was used to standardise and extract the mitogenomic data, as well as generate comparative tables for the dataset. After the removal of all unannotated mitogenomes, most hybrids between species, identical mitogenomes (we suspected species misidentification in these cases), and three species which we could not classify in terms of life history (Supplementary file S1: Text S8 and S9), the final dataset comprised 10,911 species.

Many mitogenomes had duplicated genes. As only one gene could be kept for subsequent analyses, we devised a pipeline that resolved the duplicates. The pipeline first searches for the existence of stop codons in duplicates, and then removes the one possessing them, as this implies non-functionality. If this step does not resolve the duplicates, then it compares the orthologous genes of the lowest available identical taxon in the dataset, and keeps the most conserved duplicate. This function was added to the updated version of PhyloSuite (1.2.3).

### Classification of life-history (LH) strategies

The dataset was classified according to life history into six categories: endoparasites (EndoP), ectoparasites (EctoP), parasitoids, micropredators (MP), free-living (F), and facultative parasites that exhibit intermittent parasitic and free-living life histories (EndoP/F). Endoparasites comprised parasites living inside the host’s body, which included intestines, nasal passages, etc. (so this category includes mesoparasites as well) (Supplementary file S1: Text S9). Ectoparasites attach themselves or permanently live on the outside of the host’s body. As we expected endoparasites to have a lower locomotory capacity and higher metabolic dependence on the host, we expected their evolutionary rates to be higher than those of ectoparasites. Parasitoids include organisms that are parasitic during a part of their life cycle; often parasitoids have a parasitic larva stage that eventually kills its host, followed by a free-living adult stage (Supplementary file S1: Text S9). We used parasitoids to test our hypothesis that parasites with high locomotory capacity and incomplete metabolic reliance on the host should not exhibit elevated evolutionary rates. To further confirm our predictions, we separately classified micropredators. These organisms resemble many ectoparasites in their haematophagous feeding habits, but they only visit the host for feeding, and otherwise they are free-living (e.g. mosquitoes). We hypothesised that their evolutionary rates should not differ from other free-living organisms. Finally, Strongyloididae (Nematoda) were placed in a separate category (EndoP/F) because their life cycle alternates between free-living and parasitic generations ^42,43^. We hypothesised that they should exhibit slower evolutionary rates than strictly endoparasitic nematodes. All other bilaterian species were classified as free-living.

### Classification of locomotory capacity (LC)

As we failed to find a suitable categorisation of locomotory capacity in animals, we adopted the ‘visual interaction hypothesis’, which is the best explanatory variable for variability in scaling coefficients between the mass-specific metabolic rate in marine animals: high metabolic demand follows strong selection for locomotory capacity for pursuit and evasion in visual prey/predators inhabiting well-lit oceanic waters, whereas limited visibility allows for reduced locomotory capacity, reflected in low metabolic rates ^44,45^. Such classification is of course rough, so to minimise the amount of noise interfering with analyses, we divided the dataset into three locomotory capacity categories. 1. High (H), comprising all species expected to rely on locomotion for pursuit and evasion of prey/predators. This category comprised a vast majority of species. 2. Low (L), comprising all species that have merely a rudimentary locomotory capacity (i.e. not expected to rely on locomotion for pursuit and evasion of prey/predators). This category comprised 877 species, mostly from lineages such as nematodes, flatworms, sessile molluscs and crustaceans, parasitic crustaceans, ticks, some annelids, mites, Acanthocephala, Diplura, Brachiopoda, Bryozoa, Ascidiacea, Echinodermata, Entoprocta, Gastrotricha, Gnathostomulida, Hemichordata, Kinorhyncha, Nemertea, Onychophora, Orthonectida, Phoronida, Priapulda, Tardigrada, and Xenacoelomorpha (Supplementary file S2: Worksheet A). 3. Because the distinction between the high and low locomotory capacity species is blurry in many cases, we designed a third category, Intermediate LC (I), with the aim to mop up the noise produced by these difficult-to-classify taxa, and make sure that the High and Low categories do not overlap. This category comprises species that would be expected to possess more than a rudimentary locomotory capacity, but also rely on strategies other than locomotion to evade/pursue predators/prey. Examples are certain flightless insects and Collembola, certain bathyal, abyssal and stygobitic crustacean and fish lineages, Chaetognatha, certain amphibian and reptilian lineages, some mammals, some Bivalvia and Cephalopoda, some Gastropoda, Nematomorpha, and Rotifera (Supplementary file S2: Worksheet A).

### Statistical analyses

For pairwise comparisons of branch lengths between different groups, we conducted Tukey HSD tests. As our data violated the assumption of independence due to varying levels of phylogenetic relatedness, we further addressed this problem by breaking down the dataset into pairs of categories and conducted PGLS ANOVA tests using the *nlme* package ^46^. We also conducted these tests on the overall dataset. To assess the relative impacts of different variables on branch length, we used two algorithms designed to account for the phylogenetic relatedness of data: linear fixed-effect models accounting for kinship implemented in the *lmekin* function in *coxme* ^47^, and phylogenetic multilevel Bayesian models implemented in *brms* ^48^. For both analyses, we used a matrix of phylogenetic distances extracted from the phylogenetic tree using the *ape* package, and log-transformed the branch length data to reduce the nonnormality of distribution (Supplementary file S1: Figure S12). The R^2^ parameter evaluates the proportion of the variance in the dependent variable explained by the independent variable. The R^2^ value of *lmekin* models was calculated using the “r.squaredLR” method available in the MuMIn package in R, and Bayesian R^2^ was inferred using the *bayes_r2* function of *brms* package in R. The AIC value of each *lmekin* model was calculated using the “AIC” method in R. The LRT test of *lmekin* models was conducted using the χ^2^ test following this tutorial: https://aeolister.wordpress.com/2016/07/07/likelihood-ratio-test-for-lmekin/. The outliers were removed from the overall dataset using the R “boxplot” function.

### Phylogenetic analysis

The phylogeny of Bilateria was inferred using amino acid sequences of 12 mitochondrial protein-coding genes (*atp8* was removed because it is missing from multiple lineages) of all 10,911 mitogenomes in the dataset, as well as 8 non-bilaterian Animalia outgroup species. Sequences were aligned using MAFFT ^49^, concatenated by PhyloSuite, and phylogenetic analysis conducted using FastTree 2.1.10 ^50^ under the LG evolutionary model, which accounts for the variability of evolutionary rates across sites in the matrix ^51^. The optimal model selection was performed using two approaches in ModelFinder ^52^: the AICc criterion indicated LG+G as the optimal model, whereas the BIC criterion inferred the JTT+G model. We reconstructed phylograms using both models. As the correlation between the two trees was very high (99.2%), we relied on the LG+G tree for downstream analyses. The phylum-level tree topology was constrained according to ^53^. The molecular evolution rate was defined as the root-to-tip branch lengths ^4,54^, which were extracted in two steps: the *ape* package was used to root the unrooted tree ^55^ and the patristic method in *adephylo* was used to calculate tip to root distance ^56^.

### Selection pressure analyses

To study selection pressure patterns in the dataset, we used two tools from the HYPHY suite ^57^. RELAX was used to test the hypothesis that selection pressures are relaxed in a selected subset of branches against the background branches ^58^. BUSTED was used to test the hypothesis that directional selection is associated with a certain phenotype (test branches) ^59^. As these tests require a single codon table, and codes vary across the bilaterian dataset, it had to be divided along the genetic code lines. Some of the datasets had to be further reduced to make them computationally feasible. We did this by selecting closely related test and background branches across the phylogenetic tree (details in the supplementary file S1: Text S9).

## Acknowledgements

We would like to thank Dr. Sergei L. Kosakovsky Pond for useful discussions and technical help.

## Funding sources

This work was supported by the National Natural Science Foundation of China (grant numbers 32102840 and 31872604); the Start-up Funds of Introduced Talent in Lanzhou University (561120206); the Science and Technology Project of Gansu Province (21JR7RA533). The funders had no role in study design, data collection and analysis, decision to publish, or preparation of the manuscript.

## Data availablity

All data used in this study were retrieved from the NCBI’s GenBank RefSeq database. A detailed overview of data, with GenBank accession numbers, can be found in Supplementary file S2.

## Competing interests

The authors declare no competing interest.

## Author contributions

IJ conceived the study. IJ, DZ, HZh and XL designed the analyses. DZ, IJ, CYX, and HZou were involved in the acquisition of data. DZ, CYX, IJ, and HZou performed the bioinformatic and statistical evolutionary genomic analyses. DZ developed and wrote the new code needed to conduct the analyses. IJ drafted the manuscript. DZ and GTW revised the manuscript and supervised the project.

## Notes

### Competing Interest Statement

The authors have declared no competing interest.

## References

1. Bazin, E., Glémin, S. & Galtier, N. Population Size Does Not Influence Mitochondrial Genetic Diversity in Animals. Science 312, 570–572 (2006).

2. Nabholz, B., Mauffrey, J.-F., Bazin, E., Galtier, N. & Glemin, S. Determination of Mitochondrial Genetic Diversity in Mammals. Genetics 178, 351–361 (2008).

3. Galtier, N., Jobson, R. W., Nabholz, B., Glémin, S. & Blier, P. U. Mitochondrial whims: metabolic rate, longevity and the rate of molecular evolution. Biology Letters 5, 413–416 (2009).

4. Allio, R., Donega, S., Galtier, N. & Nabholz, B. Large variation in the ratio of mitochondrial to nuclear mutation rate across animals: implications for genetic diversity and the use of mitochondrial DNA as a molecular marker. Mol Biol Evol 34, 2762–2772 (2017).

5. Jakovlić, I. et al. Drivers of interlineage variability in mitogenomic evolutionary rates in flatworms (Platyhelminthes) are multifactorial. 2022.09.11.507443 Preprint at https://doi.org/10.1101/2022.09.11.507443 (2022).

6. Dowton, M. & Austin, A. D. Increased genetic diversity in mitochondrial genes is correlated with the evolution of parasitism in the Hymenoptera. J Mol Evol 41, 958–965 (1995).

7. Shao, R., Dowton, M., Murrell, A. & Barker, S. C. Rates of Gene Rearrangement and Nucleotide Substitution Are Correlated in the Mitochondrial Genomes of Insects. Mol Biol Evol 20, 1612–1619 (2003).

8. Hassanin, A. Phylogeny of Arthropoda inferred from mitochondrial sequences: Strategies for limiting the misleading effects of multiple changes in pattern and rates of substitution. Molecular Phylogenetics and Evolution 38, 100–116 (2006).

9. Shao, R. & Barker, S. C. Mitochondrial genomes of parasitic arthropods: implications for studies of population genetics and evolution. Parasitology 134, 153–167 (2007).

10. Jakovlić, I. et al. Slow crabs - fast genomes: locomotory capacity predicts skew magnitude in crustacean mitogenomes. Molecular Ecology 30, 5488–5502 (2021).

11. Martin, P., Kaygorodova, I., Sherbakov, D. Yu. & Verheyen, E. Rapidly Evolving Lineages Impede the Resolution of Phylogenetic Relationships among Clitellata (Annelida). Molecular Phylogenetics and Evolution 15, 355–368 (2000).

12. Bernt, M. et al. A comprehensive analysis of bilaterian mitochondrial genomes and phylogeny. Molecular Phylogenetics and Evolution 69, 352–364 (2013).

13. Blaxter, M. & Koutsovoulos, G. The evolution of parasitism in Nematoda. Parasitology 142, S26–S39 (2015).

14. Gissi, C., Iannelli, F. & Pesole, G. Evolution of the mitochondrial genome of Metazoa as exemplified by comparison of congeneric species. Heredity 101, 301–320 (2008).

15. Rota-Stabelli, O. et al. Ecdysozoan Mitogenomics: Evidence for a Common Origin of the Legged Invertebrates, the Panarthropoda. Genome Biology and Evolution 2, 425–440 (2010).

16. Zou, H. et al. The complete mitochondrial genome of parasitic nematode Camallanus cotti: extreme discontinuity in the rate of mitogenomic architecture evolution within the Chromadorea class. BMC Genomics 18, 840 (2017).

17. Zhang, D. et al. Mitochondrial Architecture Rearrangements Produce Asymmetrical Nonadaptive Mutational Pressures That Subvert the Phylogenetic Reconstruction in Isopoda. Genome Biology and Evolution 11, 1797–1812 (2019).

18. Zhang, D. et al. Homoplasy or plesiomorphy? Reconstruction of the evolutionary history of mitochondrial gene order rearrangements in the subphylum Neodermata. International Journal for Parasitology 49, 819–829 (2019).

19. Castro, L. R., Austin, A. D. & Dowton, M. Contrasting Rates of Mitochondrial Molecular Evolution in Parasitic Diptera and Hymenoptera. Mol Biol Evol 19, 1100–1113 (2002).

20. Oliveira, D. C. S. G., Raychoudhury, R., Lavrov, D. V. & Werren, J. H. Rapidly Evolving Mitochondrial Genome and Directional Selection in Mitochondrial Genes in the Parasitic Wasp Nasonia (Hymenoptera: Pteromalidae). Mol Biol Evol 25, 2167–2180 (2008).

21. Ohta, T. Population size and rate of evolution. J Mol Evol 1, 305–314 (1972).

22. Page, R. D. M., Lee, P. L. M., Becher, S. A., Griffiths, R. & Clayton, D. H. A Different Tempo of Mitochondrial DNA Evolution in Birds and Their Parasitic Lice. Molecular Phylogenetics and Evolution 9, 276–293 (1998).

23. Jennings, J. B. Nutritional and respiratory pathways to parasitism exemplified in the Turbellaria. International Journal for Parasitology 27, 679–691 (1997).

24. Keeling, P. J. et al. The Reduced Genome of the Parasitic Microsporidian Enterocytozoon bieneusi Lacks Genes for Core Carbon Metabolism. Genome Biol Evol 2, 304–309 (2010).

25. Chong, R. A. & Mueller, R. L. Low Metabolic Rates in Salamanders Are Correlated with Weak Selective Constraints on Mitochondrial Genes. Evolution 67, 894–899 (2013).

26. Sun, S., Li, Q., Kong, L. & Yu, H. Limited locomotive ability relaxed selective constraints on molluscs mitochondrial genomes. Sci Rep 7, 1–8 (2017).

27. Mitterboeck, T. F. & Adamowicz, S. J. Flight loss linked to faster molecular evolution in insects. Proceedings of the Royal Society B: Biological Sciences 280, 20131128 (2013).

28. Sun, Y.-B., Shen, Y.-Y., Irwin, D. M. & Zhang, Y.-P. Evaluating the Roles of Energetic Functional Constraints on Teleost Mitochondrial-Encoded Protein Evolution. Mol Biol Evol 28, 39–44 (2011).

29. Strohm, J. H. T., Gwiazdowski, R. A. & Hanner, R. Fast fish face fewer mitochondrial mutations: Patterns of dN/dS across fish mitogenomes. Gene 572, 27–34 (2015).

30. Shen, Y.-Y., Shi, P., Sun, Y.-B. & Zhang, Y.-P. Relaxation of selective constraints on avian mitochondrial DNA following the degeneration of flight ability. Genome Res. 19, 1760– 1765 (2009).

31. Lafferty, K. D. et al. A general consumer-resource population model. Science 349, 854– 857 (2015).

32. Weinstein, S. B. & Kuris, A. M. Independent origins of parasitism in Animalia. Biology Letters 12, 20160324 (2016).

33. Huyse, T., Poulin, R. & Théron, A. Speciation in parasites: a population genetics approach. Trends in Parasitology 21, 469–475 (2005).

34. O’Leary, N. A. et al. Reference sequence (RefSeq) database at NCBI: Current status, taxonomic expansion, and functional annotation. Nucleic Acids Research 44, D733–D745 (2016).

35. Poulin, R. & Randhawa, H. S. Evolution of parasitism along convergent lines: from ecology to genomics. Parasitology 142, S6–S15 (2015).

36. Thomas, J. A., Welch, J. J., Lanfear, R. & Bromham, L. A Generation Time Effect on the Rate of Molecular Evolution in Invertebrates. Mol Biol Evol 27, 1173–1180 (2010).

37. Symonds, M. R. E. & Blomberg, S. P. A Primer on Phylogenetic Generalised Least Squares. in Modern Phylogenetic Comparative Methods and Their Application in Evolutionary Biology: Concepts and Practice (ed. Garamszegi, L. Z.) 105–130 (Springer, 2014). doi:10.1007/978-3-662-43550-2_5.

38. Thomas, J. A., Welch, J. J., Woolfit, M. & Bromham, L. There is no universal molecular clock for invertebrates, but rate variation does not scale with body size. PNAS 103, 7366–7371 (2006).

39. Nabholz, B., Glémin, S. & Galtier, N. The erratic mitochondrial clock: variations of mutation rate, not population size, affect mtDNA diversity across birds and mammals. BMC Evol Biol 9, 54 (2009).

40. Lavrov, D. V. & Pett, W. Animal mitochondrial DNA as we do not know it: Mt-Genome Organization and Evolution in Nonbilaterian Lineages. Genome Biology and Evolution 8, 2896–2913 (2016).

41. Zhang, D. et al. PhyloSuite: an integrated and scalable desktop platform for streamlined molecular sequence data management and evolutionary phylogenetics studies. Molecular Ecology Resources 20, 348–355 (2020).

42. Harvey, S. C., Gemmill, A. W., Read, A. F. & Viney, M. E. The control of morph development in the parasitic nematode Strongyloides ratti. Proceedings of the Royal Society of London. Series B: Biological Sciences 267, 2057–2063 (2000).

43. Luong, L. T. & Mathot, K. J. Facultative parasites as evolutionary stepping-stones towards parasitic lifestyles. Biology Letters 15, 20190058 (2019).

44. Childress, J. J. & Mickel, T. J. Metabolic rates of animals from the hydrothermal vents and other deep-sea habitats. Bulletin of The Biological Society of Washington 6, 249–260 (1985).

45. Seibel, B. A. & Drazen, J. C. The rate of metabolism in marine animals: environmental constraints, ecological demands and energetic opportunities. Philosophical Transactions of the Royal Society B: Biological Sciences 362, 2061–2078 (2007).

46. Pinheiro, J. et al. Package ‘nlme’. Linear and nonlinear mixed effects models, version 3, (2017).

47. Therneau, T. The lmekin function. (2018).

48. Bürkner, P.-C. brms: An R Package for Bayesian Multilevel Models Using Stan. Journal of Statistical Software 80, 1–28 (2017).

49. Katoh, K. & Standley, D. M. MAFFT multiple sequence alignment software version 7: Improvements in performance and usability. Molecular Biology and Evolution 30, 772–780 (2013).

50. Price, M. N., Dehal, P. S. & Arkin, A. P. FastTree 2 –Approximately Maximum-Likelihood Trees for Large Alignments. PLOS ONE 5, e9490 (2010).

51. Le, S. Q. & Gascuel, O. An Improved General Amino Acid Replacement Matrix. Molecular Biology and Evolution 25, 1307–1320 (2008).

52. Kalyaanamoorthy, S., Minh, B. Q., Wong, T. K. F., Von Haeseler, A. & Jermiin, L. S. ModelFinder: Fast model selection for accurate phylogenetic estimates. Nature Methods 14, 587–589 (2017).

53. Laumer, C. E. et al. Revisiting metazoan phylogeny with genomic sampling of all phyla. Proceedings of the Royal Society B: Biological Sciences 286, 20190831 (2019).

54. Lanfear, R., Thomas, J. A., Welch, J. J., Brey, T. & Bromham, L. Metabolic rate does not calibrate the molecular clock. PNAS 104, 15388–15393 (2007).

55. Paradis, E. & Schliep, K. ape 5.0: an environment for modern phylogenetics and evolutionary analyses in R. Bioinformatics 35, 526–528 (2019).

56. Jombart, T., Balloux, F. & Dray, S. adephylo: new tools for investigating the phylogenetic signal in biological traits. Bioinformatics 26, 1907–1909 (2010).

57. Kosakovsky Pond, S. L. et al. HyPhy 2.5—A Customizable Platform for Evolutionary Hypothesis Testing Using Phylogenies. Mol Biol Evol 37, 295–299 (2020).

58. Wertheim, J. O., Murrell, B., Smith, M. D., Kosakovsky Pond, S. L. & Scheffler, K. RELAX: Detecting Relaxed Selection in a Phylogenetic Framework. Mol Biol Evol 32, 820–832 (2015).

59. Murrell, B. et al. Gene-Wide Identification of Episodic Selection. Molecular Biology and Evolution 32, 1365–1371 (2015).

