## Supplementary file S1 for "Parasitism and locomotory capacity calibrate the mitogenomic evolutionary rates in Bilateria"

#### for

##### Parasitism and locomotory capacity calibrate the mitogenomic evolutionary rates in Bilateria

Ivan Jakovlić, Hong Zou, Tong Ye, Hong Zhang, Xiang Liu, Chuan-Yu Xiang, Gui-Tang Wang, Dong Zhang*

Figure S11. Branch lengths in ectoparasites classified into High and Intermediate locomotory capacity categories (EctoP_HI) vs. the free-living Insecta divided along the LC categorisation... 17

#### Supplementary Results

Table S1. The list of mitogenomes sequenced and published by our team.

| No | Species | Phylum | Life history | Publication |
|---|---|---|---|---|
| 1 | *Pseudocapillaria tomentosa* | Nematoda | Parasitic | [1] |
| 2 | *Camallanus lacustris* | Nematoda | Parasitic | [2] |
| 3 | *Clavinema parasiluri* | Nematoda | Parasitic | [2] |
| 4 | *Philometra sp.* | Nematoda | Parasitic | [2] |

| | | | | |
|---|---|---|---|---|
| 5 | *Camallanus cotti* | Nematoda | Parasitic | [3] |
| 6 | *Pingus sinensis* | Nematoda | Parasitic | [4] |
| 1 | *Cymothoa indica* | Arthropoda | Parasitic | [5] |
| 2 | *Asotana magnifica* | Arthropoda | Parasitic | [6] |
| 3 | *Tachaea chinensis* | Arthropoda | Parasitic | [7] |
| 4 | *Ichthyoxenos japonensis* | Arthropoda | Parasitic | [7] |
| 1 | *Tetraonchus monenteron* | Platyhelminthes | Parasitic | [8] |
| 2 | *Enterogyrus malmbergi* | Platyhelminthes | Parasitic | [9] |
| 3 | *Lamellodiscus spari* | Platyhelminthes | Parasitic | [10] |
| 4 | *Lepidotrema longipenis* | Platyhelminthes | Parasitic | [10] |
| 5 | *Sindiplozoon* sp. | Platyhelminthes | Parasitic | [11] |
| 6 | *Eudiplozoon* sp. | Platyhelminthes | Parasitic | [11] |
| 7 | *Paradiplozoon opsariichthydis* | Platyhelminthes | Parasitic | [11] |
| 8 | *Paratetraonchoides inermis* | Platyhelminthes | Parasitic | [12] |
| 9 | *Dactylogyrus lamellatus* | Platyhelminthes | Parasitic | [13] |
| 10 | *Thaparocleidus asoti* | Platyhelminthes | Parasitic | [14] |
| 11 | *Thaparocleidus varicus* | Platyhelminthes | Parasitic | [14] |
| 12 | *Euryhaliotrema johni* | Platyhelminthes | Parasitic | [15] |
| 13 | *Gangesia oligonchis* | Platyhelminthes | Parasitic | [16] |
| 14 | *Atractolytocestus huronensis* | Platyhelminthes | Parasitic | [17] |
| 15 | *Khawia sinensis* | Platyhelminthes | Parasitic | [17] |
| 16 | *Breviscolex orientalis* | Platyhelminthes | Parasitic | [17] |
| 17 | *Schyzocotyle acheilognathi* | Platyhelminthes | Parasitic | [17] |
| 18 | *Digramma interrupta* | Platyhelminthes | Parasitic | [18] |
| 19 | *Ligula intestinalis* | Platyhelminthes | Parasitic | [18] |
| 20 | *Gyrodactylus gurleyi* | Platyhelminthes | Parasitic | [19] |
| 21 | *Gyrodactylus kobayashii* | Platyhelminthes | Parasitic | [20] |

**Table S2. Species classification and branch length in the bilaterian dataset.** The dataset was classified according to two criteria: life history (LH) and locomotory capacity (LC). According to LH, all species were first classified into free-living and parasitic, and the latter category was then subdivided into four categories. EndoP is endoparasites, EctoP is ectoparasites, and MP is micropredators. Average and standard deviation (SD) values are shown for branch length for each category.

| **Category** | **Count** | **Branch length** | |
|---|---|---|---|
| | | **mean** | **SD** |
| **LH** | | | |
| **EctoP** | 117 | 2.011 | 0.898 |
| **EndoP** | 275 | 3.105 | 0.719 |

| | | | |
|---|---|---|---|
| **EndoP/F** | 5 | 2.385 | 0.046 |
| **Free-living** | 10264 | 1.252 | 0.274 |
| **MP** | 186 | 1.264 | 0.102 |
| **Parasitoid** | 64 | 1.423 | 0.236 |
| **LC** | | | |
| **High** | 8743 | 1.221 | 0.210 |
| **Intermediate** | 877 | 1.231 | 0.252 |
| **Low** | 1291 | 1.954 | 0.856 |

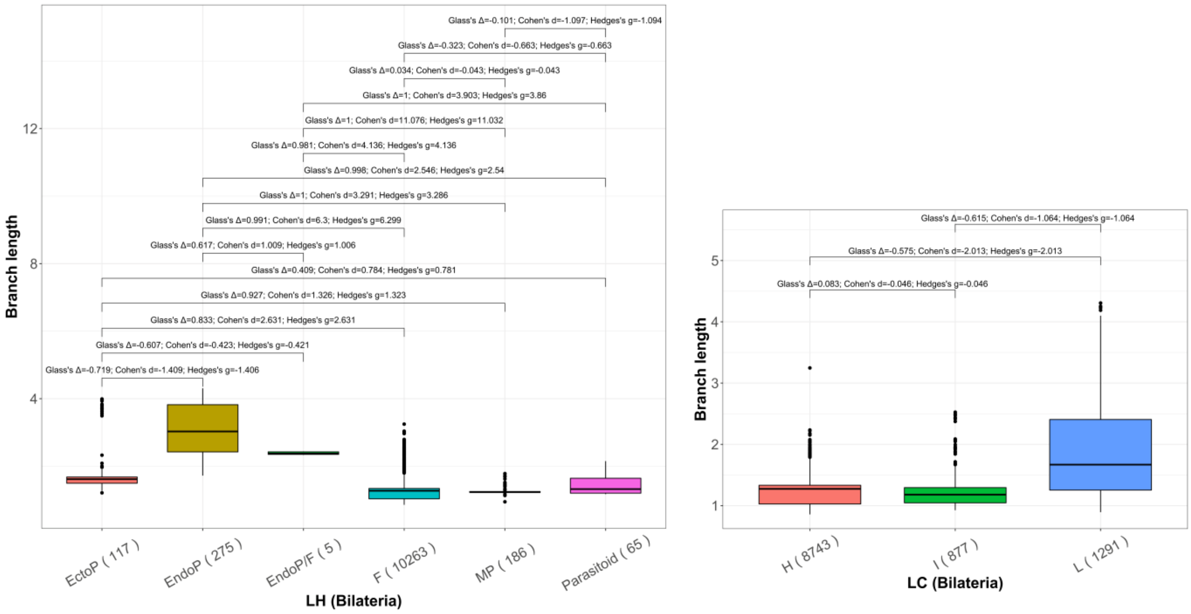

**Figure S1. The effect sizes for pairwise group comparisons.** Three different measures were used. Cohen's d should be used when the two groups have similar standard deviations and the same sample size that is >20. Glass's Δ should be used when the two groups' standard deviations are different. Hedges's g should be used when the two groups have similar standard deviations and different sample sizes, or both groups have a sample size <20 per group. In the life-history categorisation, F is free-living, EndoP is endoparasites, EctoP is ectoparasites, MP is micropredators, and EndoP/F refers to interchanging free-living and parasitic generations. In the LC categorisation, H is High, I is Intermediate, and L is Low.

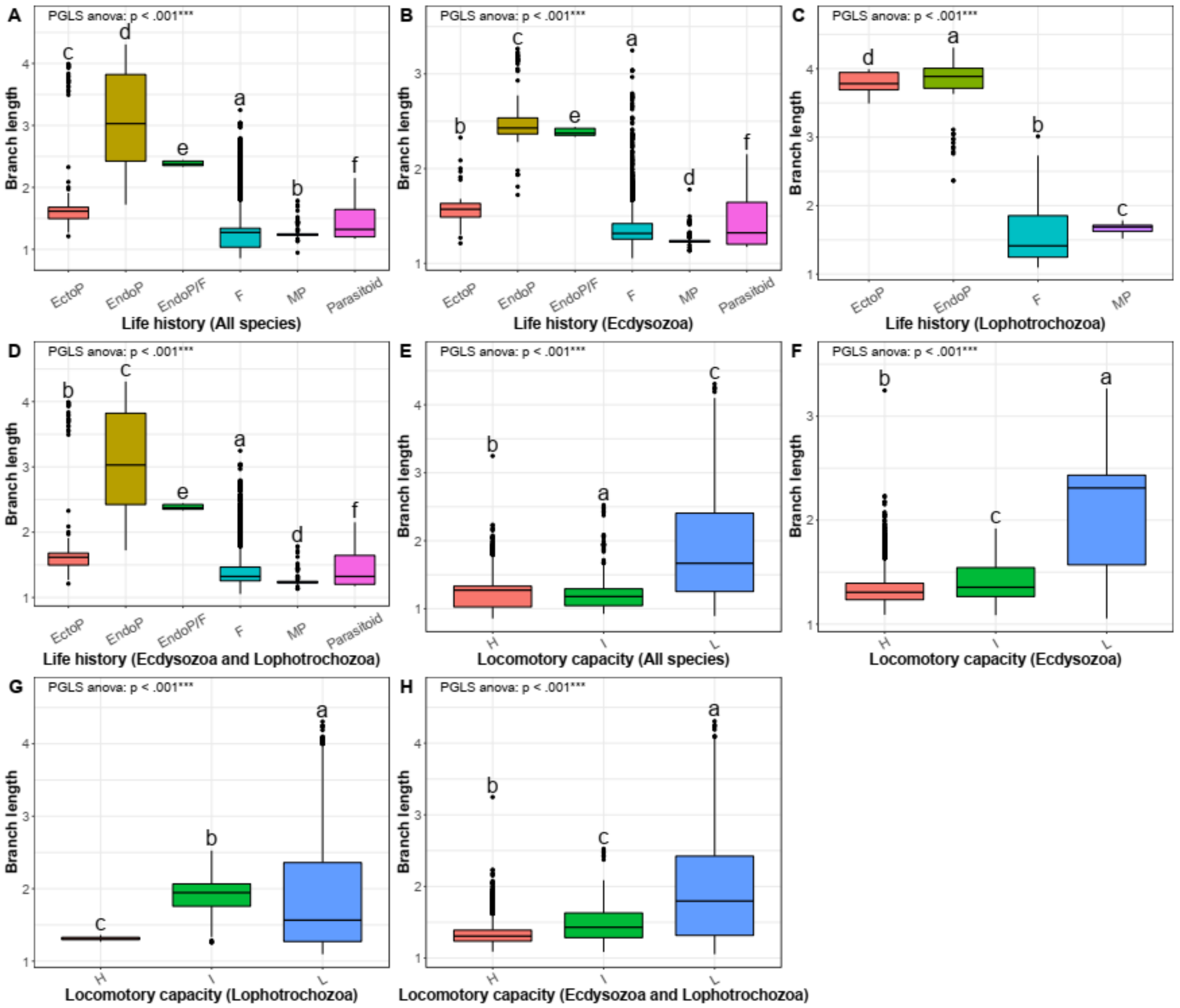

**Figure S2. PGLS ANOVA pairwise comparisons between groups and overall comparisons.** Different letters above the box plots indicate statistically significant differences. In the life-history categorisation, F is free-living, EndoP is endoparasites, EctoP is ectoparasites, and MP is micropredators. In the LC categorisation, H is High, I is Intermediate, and L is Low. Overall PGLS ANOVA results are shown in the upper left corner. Different letters above the boxplots indicate statistically significant differences ($p<0.05$).

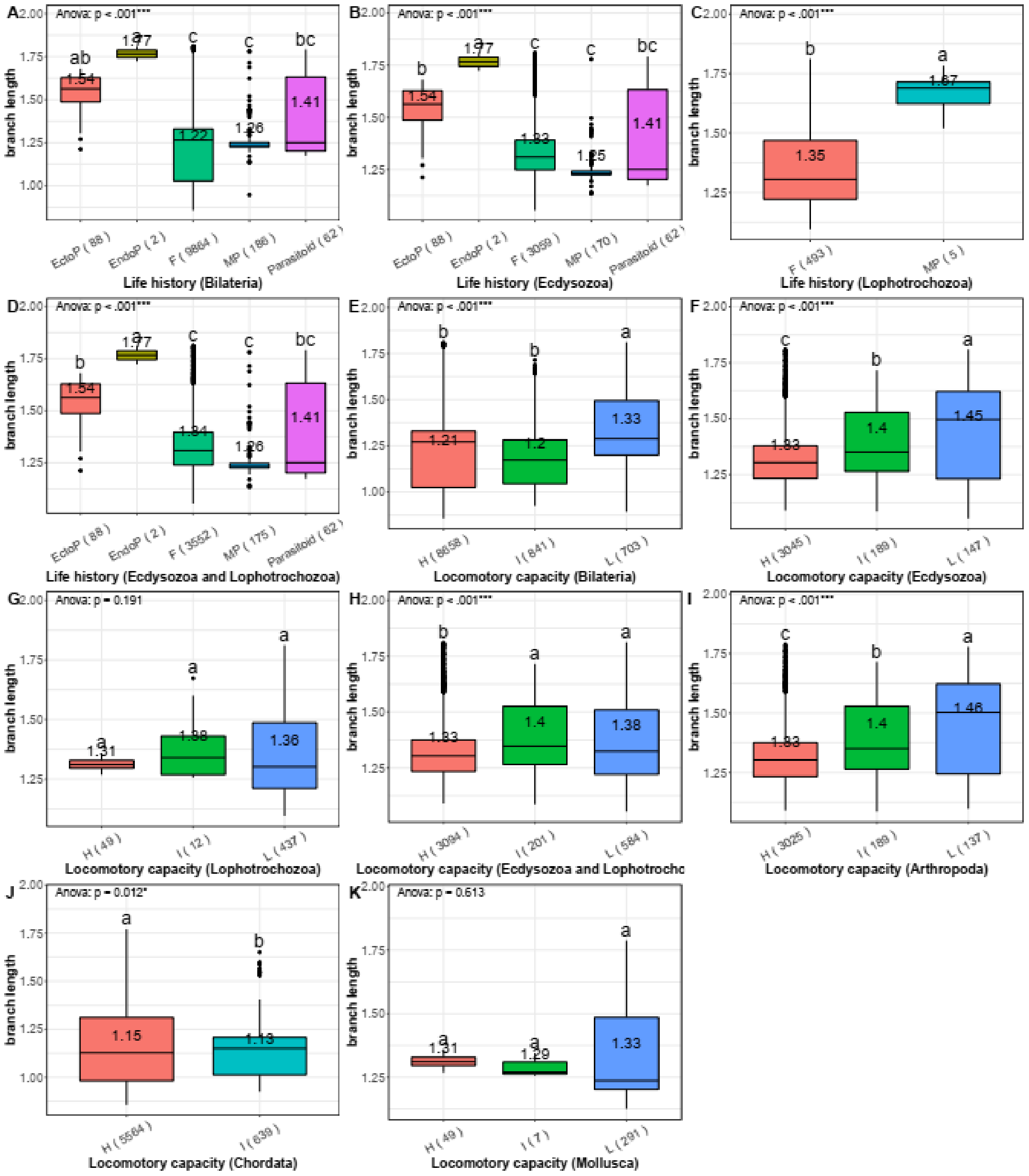

**Figure S3. Comparison of average branch lengths in different life history and locomotory capacity categories at the Bilateria level with outliers removed.** See Figure S2 for other details.

#### Text S1: Branch length comparison at the phylum-level

To assess whether the patterns observed for the entire bilaterian dataset are consistent across lower taxonomic levels, we selected taxa that comprised different life-history categories and analysed them separately. Multiple life histories were observed in four phyla, and multiple locomotory capacities in three, but they overlapped only in Arthropoda, where branch lengths mirrored the overall bilaterian pattern almost perfectly: shortest in microparasites, free-living, and parasitoids (neither of the other two differed significantly from the free-living), significantly higher in ectoparasites, and by far the highest in endoparasites (Figure S4). Similarly, branch lengths were the lowest in the high category, intermediate in the Intermediate category, and the highest in the Low locomotory capacity category (all differences were significant) (Figure S5). However, the value range was very high in the free-living category. Intriguingly, the impact of locomotory capacity outweighed the impact of all four life history categories. Apart from the ectoparasitism, all other variables had significant impacts.

In Annelida, branch lengths were significantly higher in microparasitic species, but the number of species was low (5). In the Chordata, microparasitic species did not differ significantly from the free-living, but in the LC categorisation branch lengths were almost twice as large in the Low group as in the High and Intermediate groups (almost identical between the latter two). In Nematoda, endoparasites had nonsignificantly longer branches than the free-living species. In Platyhelminthes, endoparasitic and ectoparasitic life-histories did not differ, but they both had branch lengths almost twice as long as the free-living species. The latter two phyla both had uniform LC categorisation, but in Mollusca, the Intermediate group had the longest branches, followed by the Low (1.94 vs. 1.62 respectively), and the shortest in High (all differences were significant).

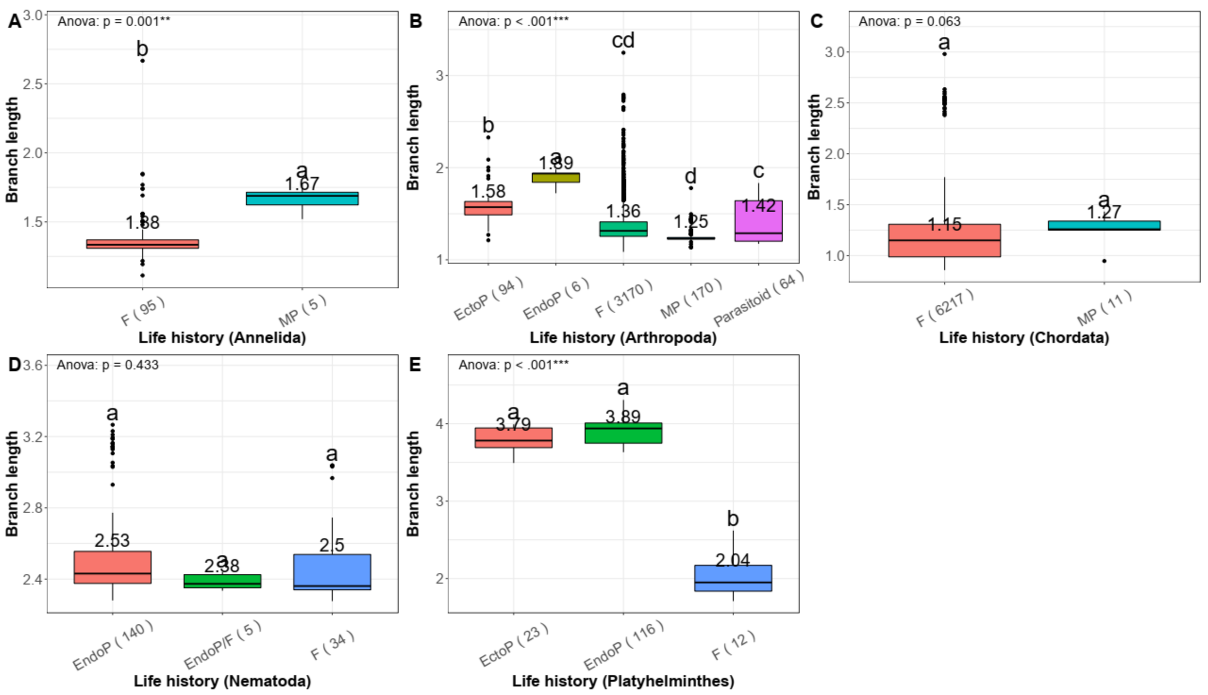

**Figure S4. Comparison of average branch lengths in different life history categories at the phylum level.** F is free-living, EndoP is endoparasites, EctoP is ectoparasites, and MP is micropredators.

Average branch length values are shown above the boxplots. PGLS ANOVA results are shown in the upper left corner. Different letters above the boxplots indicate statistically significant differences (p<0.05). The number of species included in the analysis is shown next to the category name (x-axis).

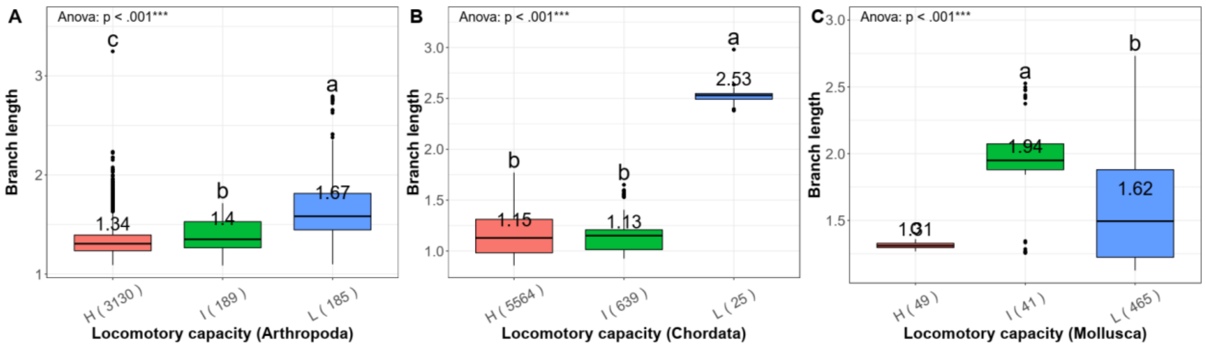

**Figure S5. Comparison of average branch lengths in different locomotory capacity categories at the phylum level.** H is High, I is Intermediate, and L is Low (locomotory capacity). Average branch length values are shown above the boxplots. PGLS ANOVA results are shown in the upper left corner. Different letters above the boxplots indicate statistically significant differences (p<0.05). The number of species included in the analysis is shown next to the category name (x-axis).

#### Text S2: Branch length comparison at the class-level

We further looked one level down, at the class level. There was more than one LC category in twelve classes, and more than one LH category in nine classes, but many analyses were weakened by a small number of samples. With regard to the life-history classification, most classes exhibited branch length patterns in agreement with the overall trend in Bilateria (Figure S6). Insecta was a class with a relatively representative sample size in most categories, so we will discuss it in more detail. Branches were the shortest in microparasites, followed by the free-living and parasitoid categories (nonsignificantly different), and the longest in ectoparasites. As regards the LC classification of insects, branches were the shortest in the High, followed by the Intermediate and the longest in the Low category. Locomotory capacity had a marginally higher impact on branch length than life history classification (both were significant, but the latter was just below the 0.05 significance threshold) (Table 2 in the main manuscript).

There were several exceptions from the bilaterian patterns in the life-history categorisation. In Clitellata and Mammalia, microparasites had longer branches than free-living, but both analyses lacked statistical power due to a low number of samples. In concordance with the phylum-level analyses, the two Nematoda classes were in disagreement with the bilaterian patterns and our hypotheses. In Chromadorea, endoparasites (2.53) had only nonsignificantly longer branches than the free-living (2.48), and in Enoplea, free-living (2.64) had nonsignificantly longer branches than endoparasitic species (2.55). Arachnida was the last apparent exception in the life history categorisation: branch lengths did not differ significantly among life history groups, but the average (and median) branch length was much lower in ectoparasites (1.58) than in free-living (1.90) species. The numbers of endoparasites (2) and microparasites (1) were too low to draw any conclusions. As this lineage was a major outlier in relation to the overall pattern, we analysed it in more detail. Two orders were outliers in terms of average branch lengths: Sarcoptiformes (2.24) and Trombidioformes (2.40). The remaining five orders had an average branch length of 1.49 (1.22 – 1.83). These two orders belong to the superorder Acariformes, which comprises most mites. The remaining mites are classified into Parasitiformes. Importantly, mites belonging to these two orders possess low locomotory capacity, similar to that exhibited by parasitic mites – ticks (Table S3). Therefore, according to our working hypothesis, there is no reason to expect faster evolution in ectoparasitic ticks with low locomotory capacity than in free-living mites with low locomotory capacity. In agreement with this, branches were significantly longer in the Low LC group than in the High LC group in Arachnida(Figure S7-C).

**Table S3. Branch lengths (brl) in the orders of Arachnida.**

| order | No. | brl | LH | LC |
|---|---|---|---|---|
| Solifugae | 2 | 1.223376 | F | H |
| Opiliones | 3 | 1.299939 | F | H |
| Scorpiones | 8 | 1.322544 | F | H |
| Ricinulei | 4 | 1.483582 | F | H |
| Ixodida | 73 | 1.573336 | EctoP | L |
| Mesostigmata | 10 | 1.667138 | EndoP+F | L |
| Araneae | | 1.831031 | F | H |
| Sarcoptiformes | 49 | 2.235278 | F | L |
| Trombidiformes | 23 | 2.395713 | F | L |

As regards locomotory capacity, High and Low categories were included only in Arachnida and three Arthropoda classes: Insecta (both discussed above), Hexanauplia (accepted as Thecostraca), and

Malacostraca). The former two classes were discussed above, and in the latter two, the distribution of average branch lengths was in agreement with our predictions: the shortest in the High, followed by the Intermediate and the highest in the Low (the analysis power was limited by only three species in Malacostraca). In other taxa, there were only two combinations: Intermediate/High and Intermediate/Low. The Intermediate group generally produced relatively noisy results, so only two of these were in agreement with our predictions (H<I<L): Chondrichthyes and Actinopteri. In Mammalia, the High and Intermediate groups were almost identical, and others (Amphibia, Bivalvia, Cephalopoda, Gastropoda, and Lepidosauria) were in disagreement with our predictions.

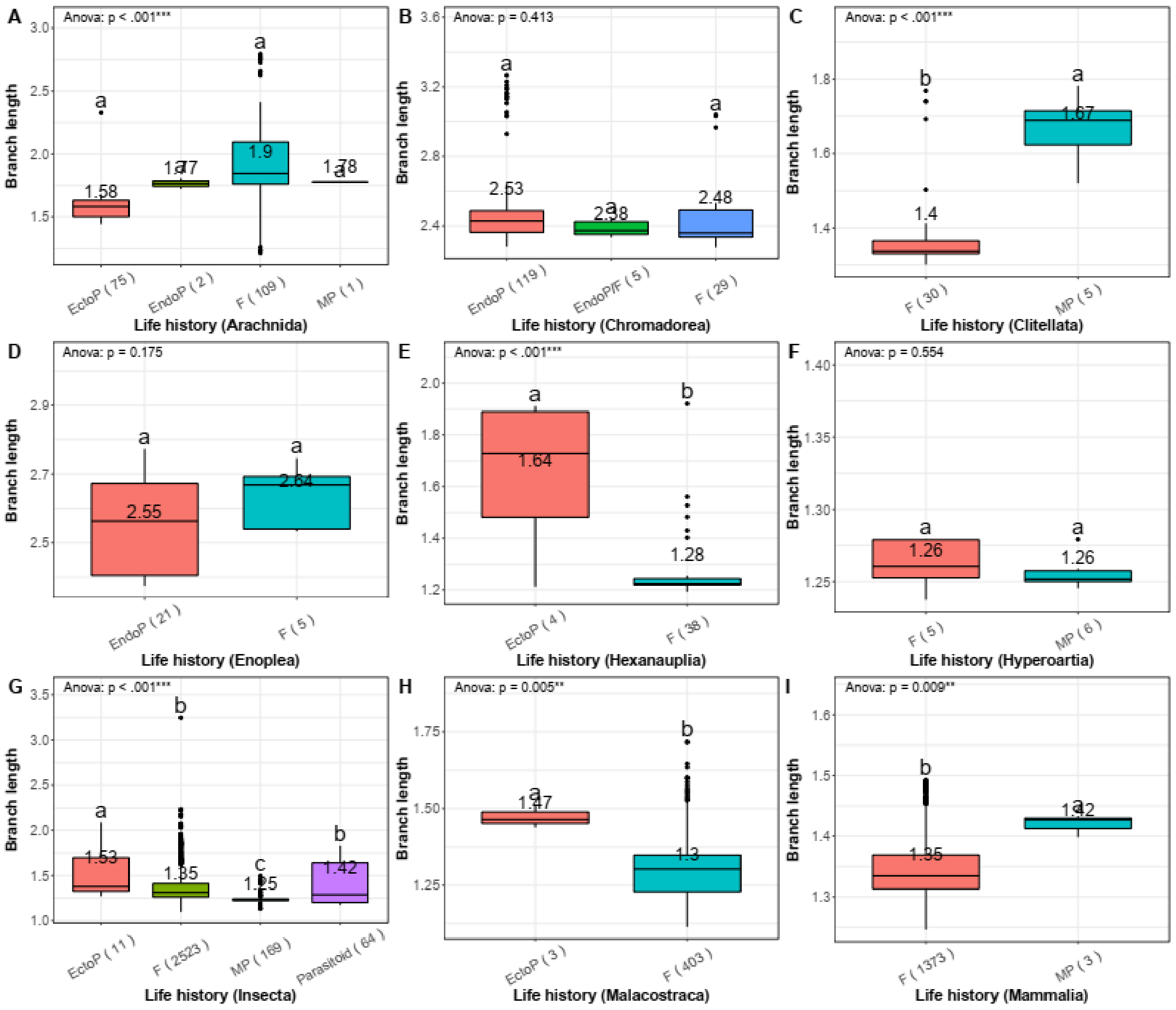

**Figure S6. Comparison of average branch lengths in different life history categories at the class level.** F is free-living, EndoP is endoparasites, EctoP is ectoparasites, and MP is micropredators. Average branch length values are shown above the boxplots. PGLS ANOVA results are shown in the upper left corner. Different letters above the boxplots indicate statistically significant differences (p<0.05). The number of species included in the analysis is shown next to the category name (x-axis). Hexanauplia is currently accepted as Thecostraca.

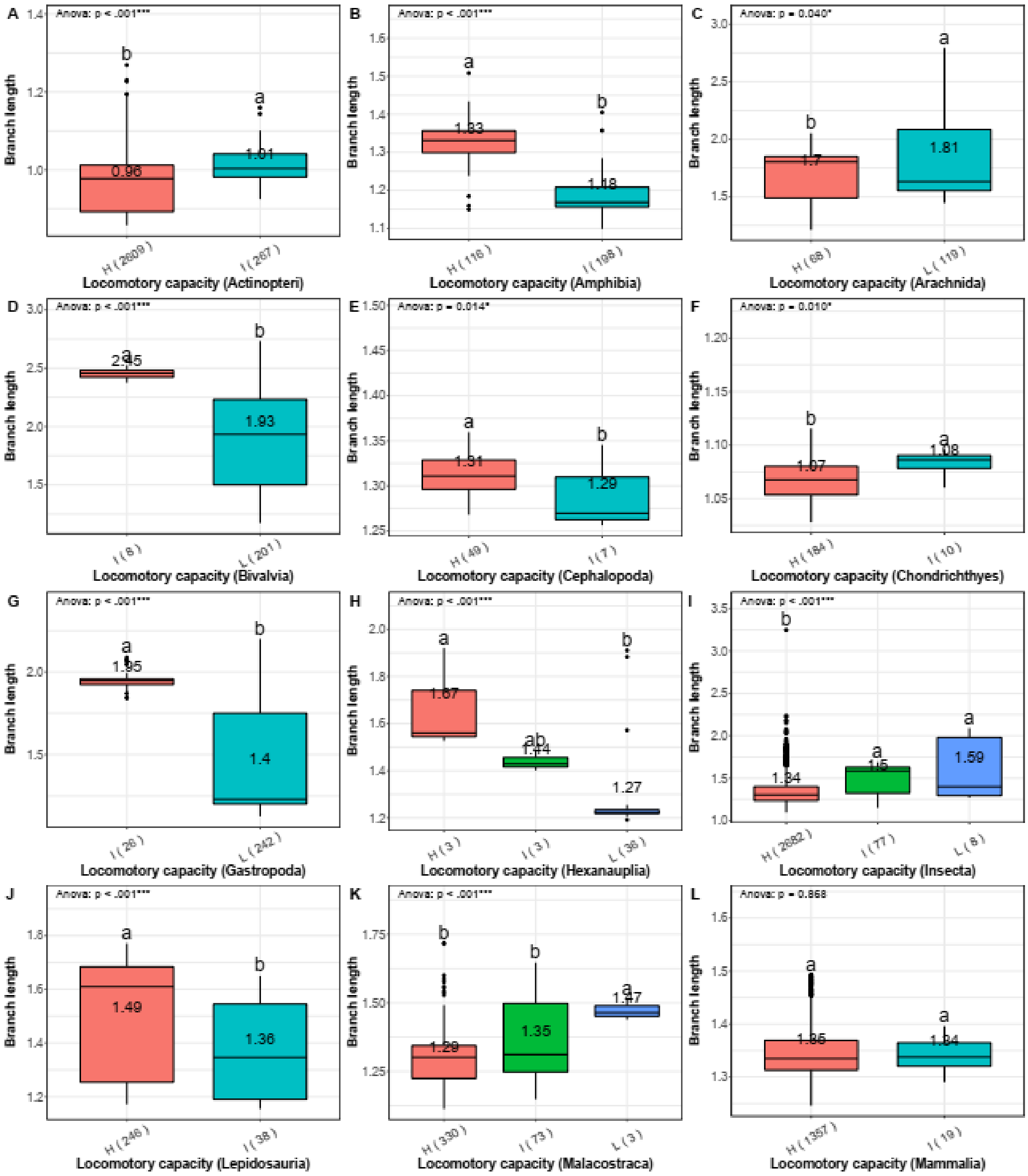

**Figure S7. Comparison of average branch lengths in different locomotory capacity categories at the class level.** H is high, I is Intermediate, and L is low (locomotory capacity). Average branch length values are shown above the boxplots. PGLS ANOVA results are shown in the upper left corner. Different letters above the boxplots indicate statistically significant differences (p<0.05). The number of species included in the analysis is shown next to the category name (x-axis). Hexanauplia is currently accepted as Thecostraca.

#### Text S3: Branch length comparison at the order-level

As regards the order level, many analyses were weakened by a low number of samples for some categories. The overall bilaterian pattern was mirrored by the Diptera (Arthropoda: Insecta): significantly longer branches in ectoparasites (only three mitogenomes) than in free-living, parasitoid and micropredator lineages, with the latter three exhibiting only marginal differences. In Rhabditida (Nematoda: Chromadorea), endoparasites exhibited much higher average values than free-living and Endop/F species, but differences were nonsignificant, possibly due to a number of outliers in the free-living group and only five samples in the endoparasitic group. There were no significant differences between the free-living and parasitoids in Hymenoptera, and free-living and microparasites in Hemiptera, Petromyzontiformes, and Hirudinida. In Chiroptera, microparasites (only three species) had significantly longer branches.

As regards the locomotory capacity, most orders merely had High and Intermediate categories. In most cases, differences were small and nonsignificant. Anura was the only lineage with a relatively large and significant difference: the High group had longer branches than the Intermediate group.

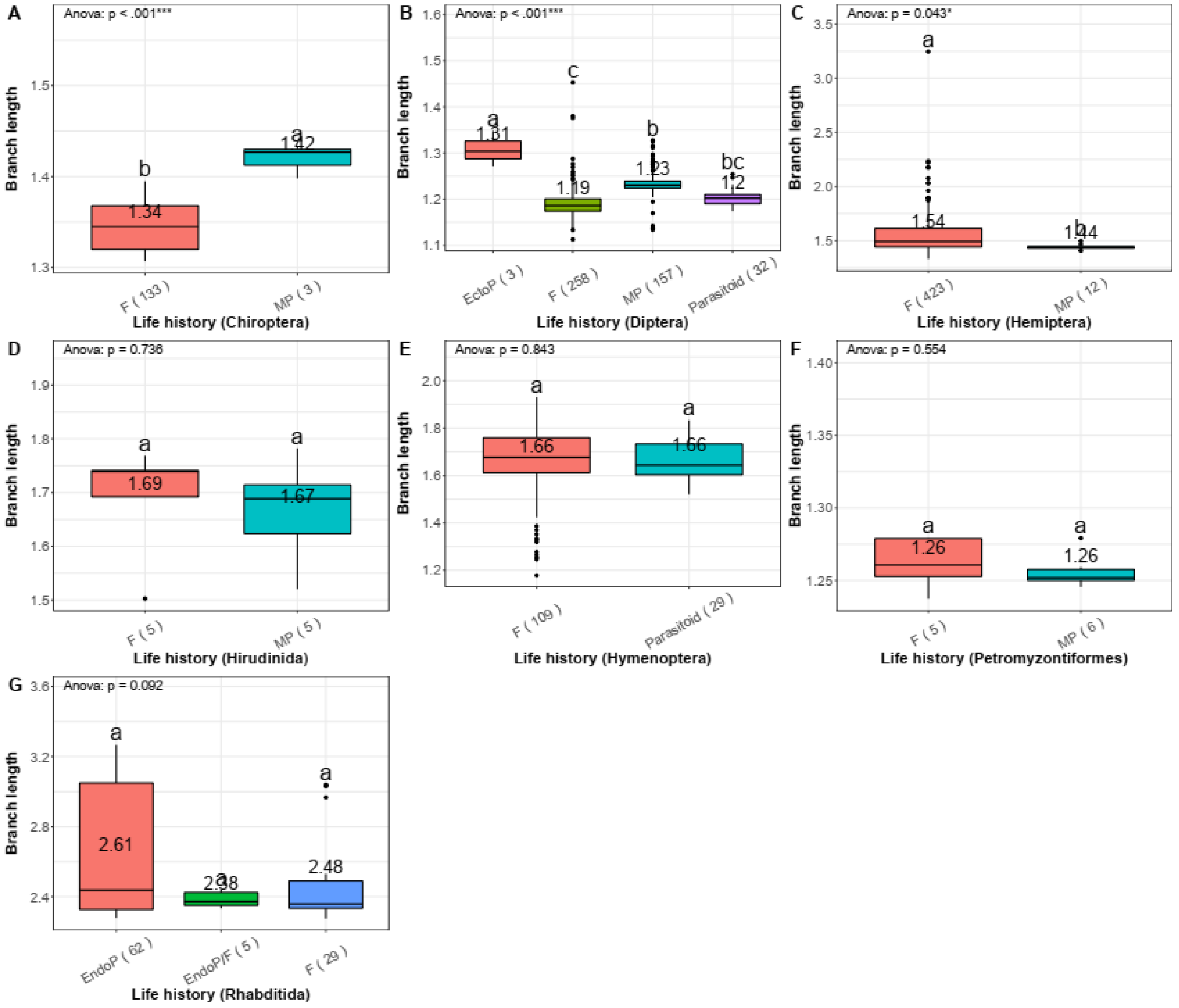

**Figure S8. The comparison of average branch lengths in different life history categories at the order level.** F is free-living, EndoP is endoparasites, EctoP is ectoparasites, and MP is micropredators. Average branch length values are shown above the boxplots. PGLS ANOVA results are shown in the upper left corner. Different letters above the boxplots indicate statistically significant differences (p<0.05). The number of species included in the analysis is shown next to the category name (x-axis).

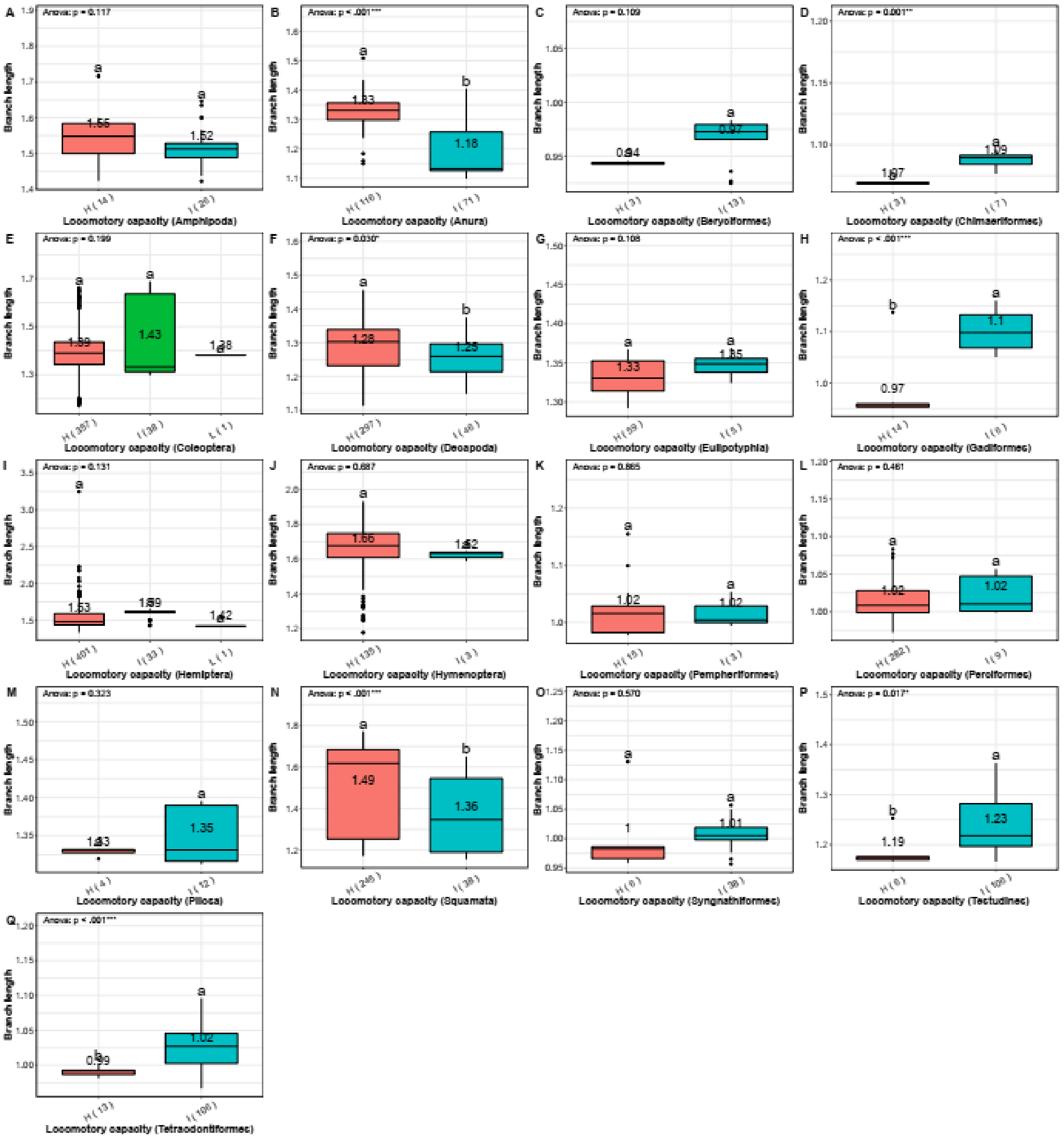

**Figure S9. The comparison of average branch lengths in different locomotory capacity categories at the order level.** H is High, I is Intermediate, and L is Low (locomotory capacity). Average branch length values are shown above the boxplots. PGLS ANOVA results are shown in the upper left corner. Different letters above the boxplots indicate statistically significant differences (p<0.05). The number of species included in the analysis is shown next to the category name (x-axis).

#### Text S4: Single gene results

We further assessed whether single-gene topologies may produce different results by constructing topology-constrained single-gene trees for all 12 PCGs, extracting branch lengths and assessing correlation with the main tree (produced using concatenated genes). Branch lengths of single-gene trees generally had a high correlation of over 80%, and most of them exhibited values over 68%. The only exceptions were *nad2*, *nad4L*, and *nad6*, which had correlations of 42 to 46% (Table S4). As *nad2* is the only relatively large gene among the three outlier genes (≈1000 bp), we conducted pairwise comparative analyses using only the *nad2* branch length data. In the main aspects, the results were congruent with the overall dataset (Supplementary file S1: Figure S10).

**Table S4. Correlation between branch lengths in single-gene phylogenetic trees and concatenated dataset (12PCGs) phylogenetic trees inferred using two different evolution models (JTT and LG).**

| tree | *atp6* | *cox1* | *cox2* | *cox3* | *cytb* | *nad1* | *nad2* | *nad3* | *nad4l* | *nad4* | *nad5* | *nad6* | 12PCGs_JTT | 12PCGs_LG |
|---|---|---|---|---|---|---|---|---|---|---|---|---|---|---|
| ***atp6*** | 1 | 0.8004 | 0.8254 | 0.8117 | 0.7411 | 0.7798 | 0.3430 | 0.7345 | 0.3492 | 0.7919 | 0.7658 | 0.3481 | 0.8815 | 0.8829 |
| ***cox1*** | 0.8004 | 1 | 0.7657 | 0.9254 | 0.6424 | 0.6514 | 0.1527 | 0.7020 | 0.3070 | 0.6913 | 0.7465 | 0.1104 | 0.8843 | 0.9025 |
| ***cox2*** | 0.8254 | 0.7657 | 1 | 0.8342 | 0.8851 | 0.7905 | 0.6016 | 0.5804 | 0.5213 | 0.8171 | 0.5610 | 0.5482 | 0.9233 | 0.9223 |
| ***cox3*** | 0.8117 | 0.9254 | 0.8342 | 1 | 0.7619 | 0.7359 | 0.3365 | 0.6626 | 0.3539 | 0.7764 | 0.6766 | 0.2927 | 0.9372 | 0.9436 |
| ***cytb*** | 0.7411 | 0.6424 | 0.8851 | 0.7619 | 1 | 0.7475 | 0.7018 | 0.4541 | 0.5012 | 0.7571 | 0.3738 | 0.6481 | 0.8794 | 0.8697 |
| ***nad1*** | 0.7798 | 0.6514 | 0.7905 | 0.7359 | 0.7475 | 1 | 0.5332 | 0.6270 | 0.3288 | 0.9013 | 0.6783 | 0.6154 | 0.8551 | 0.8249 |
| ***nad2*** | 0.3430 | 0.1527 | 0.6016 | 0.3365 | 0.7018 | 0.5332 | 1 | 0.0611 | 0.4957 | 0.5089 | -0.0201 | 0.7764 | 0.4864 | 0.4618 |
| ***nad3*** | 0.7347 | 0.7020 | 0.5804 | 0.6626 | 0.4541 | 0.6270 | 0.0611 | 1 | 0.3758 | 0.6459 | 0.8102 | 0.0964 | 0.6800 | 0.7022 |
| ***nad4L*** | 0.3492 | 0.3070 | 0.5213 | 0.3539 | 0.5012 | 0.3288 | 0.4957 | 0.3758 | 1 | 0.3558 | 0.1503 | 0.3706 | 0.4137 | 0.4507 |
| ***nad4*** | 0.7919 | 0.6913 | 0.8171 | 0.7764 | 0.7571 | 0.9013 | 0.5089 | 0.6459 | 0.3558 | 1 | 0.6923 | 0.5658 | 0.8779 | 0.8582 |
| ***nad5*** | 0.7658 | 0.7465 | 0.5610 | 0.6766 | 0.3738 | 0.6783 | -0.0201 | 0.8102 | 0.1503 | 0.692324 | 1 | 0.0581 | 0.6842 | 0.6855 |
| ***nad6*** | 0.3481 | 0.1104 | 0.5482 | 0.2927 | 0.6481 | 0.6154 | 0.7764 | 0.0964 | 0.3706 | 0.5658 | 0.0581 | 1 | 0.4652 | 0.4243 |
| **12PCGs_JTT** | 0.8815 | 0.8843 | 0.9233 | 0.9372 | 0.8794 | 0.8551 | 0.4864 | 0.6800 | 0.4137 | 0.8779 | 0.6842 | 0.4652 | 1 | 0.9925 |
| **12PCGs_LG** | 0.8829 | 0.9025 | 0.9223 | 0.9436 | 0.8697 | 0.8249 | 0.4618 | 0.7022 | 0.4507 | 0.8581 | 0.6855 | 0.4243 | 0.9925 | 1 |

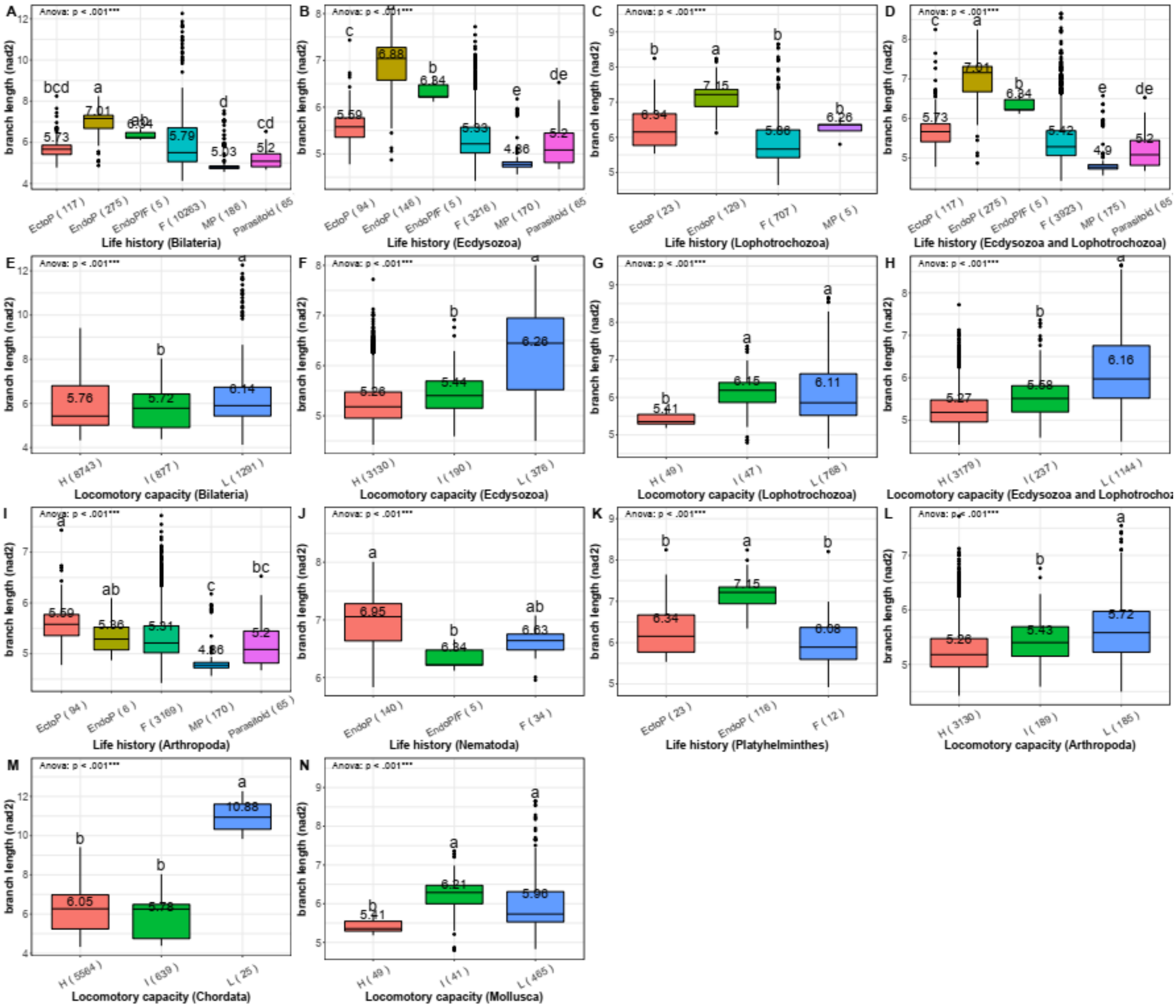

**Figure S10. nad2 tree branch length comparison between different life history categories and locomotory capacity categories.** In the life-history categorisation, F is free-living, EndoP is endoparasitic, EctoP is ectoparasitic, and MP is micropredatory. In the LC categorisation, H is high, I is Intermediate, and L is Low. Average branch length values are shown above the boxplots. PGLS ANOVA results are shown in the upper left corner. Different letters above the boxplots indicate statistically significant differences (p<0.05). The number of species included in the analysis is shown next to the category name (x-axis).

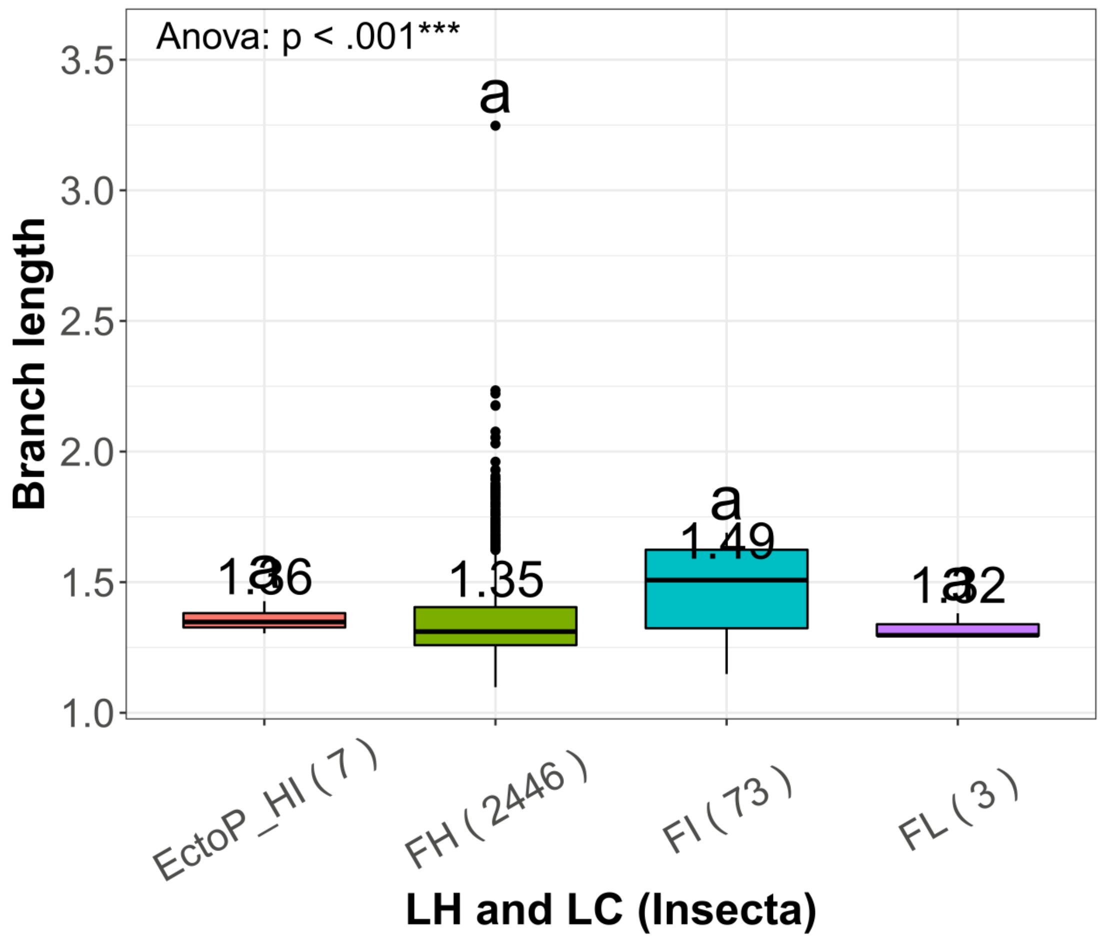

**Figure S11. Branch lengths in ectoparasites classified into High and Intermediate locomotory capacity categories (EctoP_HI) vs. the free-living Insecta divided along the LC categorisation.** FH: free-living lineages with high LC, FI: free-living with Intermediate LC, FL: free-living with low LC. Average branch length values are shown above the boxplots. PGLS ANOVA results are shown in the upper left corner. Different letters above the boxplots indicate statistically significant differences (p<0.05). The number of species included in the analysis is shown next to the category name (x-axis).

**Table S5. Phylogeny-corrected lmekin and brms regression analyses with branch length as the dependent variable and dataset divided along different life history lines as the independent variable.** $R^2$ evaluates how much a variable can explain the variance of the dependent variable (calculated via univariate regression). $aR^2$ is the adjusted $R^2$ value. F is free-living, EndoP is endoparasites, EctoP is ectoparasites, and MP is micropredators. In the *brms* analyses, every parameter is summarized using the mean (Est), standard deviation (Error) of the posterior distribution, as well as two-sided 95% Credible intervals (l and u) based on quantiles.

| **Dataset** | **LMEKIN** | | **BRMS** | | | | | | | |
|---|---|---|---|---|---|---|---|---|---|---|
| | **$R^2$** | **$aR^2$** | **Est** | **Error** | **l** | **u** | **$R^2$** | **$R^2$ Err** | **Q2.5** | **Q97.5** |
| EndoP vs. EctoP vs. free-living (F+MP) vs. parasitoid | 0.3489 | 0.3706 | 1.93 | 0.09 | 1.76 | 2.09 | 0.3474 | 0.0060 | 0.3358 | 0.3587 |
| EndoP vs. EctoP vs. free-living (F+MP+parasitoid) | 0.3470 | 0.3686 | 1.93 | 0.09 | 1.76 | 2.11 | 0.3456 | 0.0060 | 0.3338 | 0.3572 |
| Parasitic (EndoP+EctoP) vs free-living (F+MP) vs. parasitoid | 0.3215 | 0.3415 | 3.03 | 0.04 | 2.95 | 3.11 | 0.3195 | 0.0060 | 0.3079 | 0.3313 |
| Parasitic (EndoP+EctoP) vs. free-living (F+MP+parasitoid) | 0.3195 | 0.3394 | 3.03 | 0.04 | 2.95 | 3.11 | 0.3177 | 0.0060 | 0.3060 | 0.3296 |

**Table S6. BRMS multilevel regression analyses with branch length as the dependent variable.** Every parameter is summarized using the mean (Est.), standard deviation (Error) of the posterior distribution, as well as two-sided 95% Credible Intervals (l-95% and u-95%) based on quantiles. $R^2$ evaluates how much a variable can explain the variance of the dependent variable (calculated via univariate regression). The random effect was not included in any of the analyses. Ind var = Independent variable, ncp = not corrected for phylogeny. LH×LC tests the interaction between the two variables.

| Dataset | Ind var | Est | Est Error | l-95% | u-95% | $R^2$ | $R^2$ Error | Q2.5 | Q97.5 |
|---|---|---|---|---|---|---|---|---|---|
| Bilateria | LH | 1.92 | 0.09 | 1.74 | 2.10 | 0.3498 | 0.0058 | 0.3385 | 0.361114 |
| Bilateria | LC | 0.02 | 0.03 | -0.03 | 0.08 | 0.2654 | 0.0062 | 0.2535 | 0.277732 |
| Bilateria | LH+LC | 1.87 | 0.08 | 1.71 | 2.03 | 0.4160 | 0.0052 | 0.4058 | 0.425927 |
| Bilateria | LH+LC(ncp) | 1.87 | 0.08 | 1.71 | 2.05 | 0.4161 | 0.0055 | 0.4045 | 0.426748 |
| Bilateria | Null model | N/A | N/A | N/A | N/A | 2.00E-30 | 4.38E-30 | 1.27E-34 | 1.37E-29 |
| Ecdysozoa | LH | 2.48 | 0.10 | 2.28 | 2.68 | 0.4378 | 0.0091 | 0.4197 | 0.454801 |
| Ecdysozoa | LC | 0.24 | 0.06 | 0.13 | 0.35 | 0.4156 | 0.0095 | 0.3969 | 0.434494 |
| Lophotrochozoa | LH | -0.01 | 0.13 | -0.26 | 0.23 | 0.7025 | 0.0096 | 0.6822 | 0.719847 |
| Lophotrochozoa | LC | 0.86 | 0.21 | 0.45 | 1.26 | 0.0386 | 0.0126 | 0.0171 | 0.065638 |
| Ecdysozoa+Lophotrochozoa | LH | 1.83 | 0.08 | 1.68 | 1.99 | 0.5278 | 0.0069 | 0.5141 | 0.541093 |
| Ecdysozoa+Lophotrochozoa | LC | 0.38 | 0.05 | 0.28 | 0.49 | 0.3152 | 0.0092 | 0.2968 | 0.333118 |
| Bilateria | LH×LC | -320.05 | 135.54 | -603.60 | -92.47 | 0.4164 | 0.0056 | 0.4053 | 0.426969 |
| F | Low LC | 0.03 | 0.03 | -0.03 | 0.10 | 0.1077 | 0.0056 | 0.0965 | 0.118392 |
| F | Null-model | N/A | N/A | N/A | N/A | 1.89E-30 | 3.94E-30 | 1.75E-34 | 1.33E-29 |
| Low LC | LH | 1.15 | 0.08 | 0.98 | 1.31 | 0.4787 | 0.0152 | 0.4483 | 0.507245 |
| Low LC | Null-model | N/A | N/A | N/A | N/A | 2.68E-31 | 6.15E-31 | 2.08E-35 | 1.93E-30 |

**Text S5: Selection analyses**

The bilaterian dataset comprised three genetic codes. Among the genetic code table 5 lineages, we selected all endoparasites, ectoparasites, parasitoids, and a subset of Low LC species, and compared each to a different selection of free-living lineages. We further tested all Nematoda (code table 5; EndoP vs. F), and Platyhelminthes (code table 9; EndoP, EctoP and F) separately. As regards the LC categorisation, there were 3187 High, 1006 Low, and 238 Intermediate species. With Low LC lineages selected as the test branches, and Intermediate and High LC as reference branches, the test branches had a higher ω than reference branches (0.0442 vs. 0.0405). The purifying selection pressures were generally relaxed in endoparasites, parasitoids, and Low LC group, but not in ectoparasites, in comparison to the free-living (Table S7). However, the results were rather noisy and inconsistent. For example, the selection was intensified in endoparasites compared to ectoparasites in the code table 5 dataset. In flatworms, the selection was nonsignificantly relaxed in endoparasites vs. the free-living, intensified in ectoparasites compared to the free-living, and relaxed in endoparasites vs. ectoparasites. There were signals of significant directional selection in both parasitic and free-living lineages in most cases. Exceptions were parasitoids (no selection in the selected free-living background), and flatworms, where results were again contradictory, but in general there was no directional selection in the free-living lineages (Table S7). However, the flatworm dataset had very few free-living species (11). Previously, we used a larger flatworm dataset, comprising all mitogenomes available in the NCBI's Nucleotide database (30 free-living species), and found that selection was highly significantly relaxed in endoparasitic flatworms ($p < 0.0001$; Supplementary file S1: Table S8) [21]. On this basis, we can infer that the above outlier result is almost certainly an artefact caused by the small number of free-living species. We can conclude that a proportion of the noise certainly stems from the fact that the Bilaterian dataset had to be strongly reduced due to it comprising different genetic codes and to make it computationally feasible (more in Methods). These analyses may have been further confounded by the violations of assumptions of the parameter-rich algorithms employed, artefactual relationships in phylogenetic reconstructions, as well as high levels of mutational saturation of data comprising such a wide phylogenetic scope of species.

**Table S7. RELAX and BUSTED selection pressure tests. RELAX tests the hypothesis that purifying selection pressure is relaxed in the test branches (left) vs. the background branches (right).** BUSTED tests the hypothesis that episodic diversifying positive selection occurred on test and/or background branches. In brackets next to group names are the numbers of species included in the dataset. K is a measure of selection pressure, where values <1 indicate purifying selection, > directional selection, and 1 indicates neutral selection. LRT is the likelihood ratio test.

| | | **RELAX** | | | | **BUSTED** | |
|---|---|---|---|---|---|---|---|
| **Dataset** | **Compared groups** | **Selection** | **K** | **LRT** | **p** | **Test** | **Background** |
| Code table 5 | EndoP (159) vs. F (61) | relaxation | 0.641 | 390.792 | <0.0001 | <0.0001 | <0.0001 |
| | EndoP (159) vs. EctoP (94) | intensification | 1.334 | 575.674 | <0.0001 | <0.0001 | <0.0001 |
| | EctoP (94) vs. F (84) | neutral | 0.999 | 0.018 | 0.8926 | <0.0001 | <0.0001 |
| EctoP | Low LC (87) vs. High LC (6) | intensification | 1.166 | 45.776 | <0.0001 | <0.0001 | <0.0001 |
| Parasitoid | Parasitoid (65) vs. F (65) | relaxation | 0.784 | 354.466 | <0.0001 | <0.0001 | 0.5000 |
| Nematoda | EndoP (140) vs. F (34) | relaxation | 0.779 | 501.181 | <0.0001 | <0.0001 | <0.0001 |
| Platyhelminthes | EndoP (114) vs. F (11) | neutral | 0.981 | 2.651 | 0.1035 | <0.0001 | 0.5000 |
| | EctoP (23) vs. F (11) | intensification | 1.060 | 56.806 | <0.0001 | 0.5000 | 0.5000 |
| | EndoP (114) vs. EctoP (23) | relaxation | 0.834 | 206.531 | <0.0001 | 0.5000 | <0.0001 |
| LC table 5 | Low LC (210) vs. High LC (233) | relaxation | 0.951 | 26.403 | <0.0001 | <0.0001 | <0.0001 |

**Table S8. Selection tests conducted on mitogenomes of all Platyhelminthes species available in the NCBI's Nucleotide database.** See Table S7 for other details.

| RELAX | Dataset | Selection | K | LRT | P |
|---|---|---|---|---|---|
| | EndoP (152) vs. F (30) | relaxation | 0.839 | 118.119 | <0.0001 |
| | EctoP (41) vs. F (30) | | | | error |
| | EndoP (152) vs. EctoP (41) | relaxation | 0.901 | 122.744 | <0.0001 |
| **BUSTED** | **Dataset** | **P** | **P** | | |
| | EndoP (152) vs. F (30) | <0.0001 | 0.5000 | | |
| | EctoP (41) vs. F (30) | 0.1146 | 0.5000 | | |
| | EndoP (152) vs. EctoP (41) | 0.1954 | <0.0001 | | |

### Supplementary discussion

#### Text S6: Additional discussion of potentially confounding variables

Many variables have been associated with mitogenomic evolution, and the parsing of their impacts is further complicated by their interrelatedness [22]. It is beyond the scope of this study to discuss all of them in detail, but we will try to address the major ones succinctly.

Effects of some variables appear to be lineage-specific: in vertebrates, body size/mass and metabolic rate calibrate the (mitochondrial and nuclear) molecular clock (smaller animals = faster evolution) [23–25], but there is no evidence for this in invertebrates [22,26,27]. Indeed, parasitic nematodes are on average much larger than free-living nematodes, and some parasitic flatworms are much larger than free-living ones. Generation time appears to have effects in both vertebrates and invertebrates: organisms with shorter generation times have a greater number of germ-cell divisions per year, and thus more replication errors per time unit, leading to faster mutation rates [23,28]. A closely related variable, longevity (or lifespan), was also associated with mitogenomic evolution. The hypothesis proposes that long-lived animals may have adapted to an increased lifespan by evolving macromolecular components more resistant to oxidative damage, thus reducing their evolutionary rates [29–31]. This hypothesis has been thoroughly tested in mammals, and several studies also found support for it in invertebrates [28,30,32,33]. However, multiple studies reported that there are exceptions from the expected negative relationship between longevity and/or generation time and molecular clock rate [21,29,34–37]. While parasitic lineages do have shorter generation times than their vertebrate hosts [38], this may not be a rule in invertebrates [39]. For example, many parasitic flatworms and nematodes have complex life cycles that require multiple hosts and maturation stages, which results in rather long generation times [38]. As many small invertebrates have very short generation times [28,40], it is unclear whether there is a significant difference in generation times between closely related parasitic and free-living invertebrate lineages. It would be interesting to study the impact of this variable on mitogenomic evolution in parasitic lineages, but the limited availability of data may be a confounding factor.

The effective population size ($N_e$) is positively correlated with the strength of purifying selection, so species with lower $N_e$ should exhibit higher rates of mitochondrial evolution [41]. This also implies that the speciation rate may be positively correlated with the evolutionary rate via the founder effect [42]. The evidence for the putative correlation between the $N_e$ and mitochondrial evolution remains elusive: a case study found a negative correlation [43], but several studies of much larger datasets found weak evidence [44], or no evidence at all [21,22,45]. A study argued that this may be due to large differences in mutation rates [36], so the impact of $N_e$ remains debatable. It should be noted that molecular *Ne* estimates are sensitive to a number of methodological and life history parameters [46,47], so *Ne* inference can be rather difficult and error-prone. Most importantly, the *Ne* is highly likely to fluctuate strongly throughout the evolutionary history of any lineage [48], so the current *Ne* values may be completely decoupled from the root to tip branch lengths of the lineage.

Several studies found support for the hypothesis that thermic habitat affects mitochondrial evolution [49,50] and that evolutionary rates differ between endotherms and ectotherms [34]. The 'functional constraints' hypothesis proposes that variations in the thermic environment may restrict physiologically acceptable amino acid substitutions, which implies that thermally stable endotherms should have a higher rate of sequence evolution than thermally variable ectotherms [34,51]. However, several studies found that the impact of thermic habitat is very weak or inconsistent [21,34,51,52].

Mutation pressure is presumed to be positively correlated with the metabolic rate: a high metabolic rate increases the overall time that the mitochondrial H-strand spends in the mutagenic single-strand state (replication and transcription) and increases the production of mutagenic reactive oxygen species [23,27,34,53–55]. However, studies failed to find evidence for the association of metabolic and mutational rates in isopods [37], crustaceans [22], and across Metazoa [27].

Another important variable may be variability in mtDNA replication and repair mechanisms: lower fidelity = higher evolution [28,56–59]. However, multiple independent origins of parasitism in Bilateria [60] suggest that it is statistically highly unlikely that random variations in this variable may explain the observed patterns. In other words, variation in mitogenomic replication and repair mechanisms in bilaterians is probably also tightly associated with the variability in purifying selection pressures, which are at least partially driven by the selection for mitogenomic metabolic efficiency. For example, it has been proposed previously that elevated evolutionary rates in nematodes may be attributable to the loss of a mitochondrial replicase subunit [59]. As the entire phylum exhibits a limited locomotory capacity, this would explain why such a loss was not strongly selected against, as it would have been in a highly locomotory lineage.

Several previous studies attributed elevated evolutionary rates in parasites to directional selection and genetic draft (comprising selective sweeps and compensation-draft feedback) [28,44,61–64]. However, if the high evolutionary rate in parasites is driven by the arms race between parasites and hosts, then we would expect host species to exhibit equally fast evolutionary rates, or otherwise parasites would be winning the race. Therefore, we hypothesised that both parasitic and free-living lineages should be undergoing directional selection, but parasitic lineages should exhibit relaxed purifying selection pressures compared to the free-living lineages. We found some support for this hypothesis, but the results were relatively noisy, which may have been caused by a plethora of factors (more in Text S5). It is possible that relaxed purifying selection pressures allow an increased number of mutations to accumulate in parasitic organisms, which in turn provides a richer material for directional selection to work on compared to lineages evolving under strict purifying selection pressures. Intriguingly, rates of adaptive substitution are substantially higher in invertebrates than in vertebrates [45,65], which may also be a reflection of the above mechanism.

Finally, a variable that received limited scientific attention so far is the reduction of metabolic and genomic complexity in some parasites, putatively in combination with high metabolic dependence on the host [66–68]. Herein, we hypothesised that this may allow a degradation of the mitogenomic energy production efficiency, i.e. relaxed purifying selection pressures in some parasitic lineages. We provide additional discussion of some major variables in Supplementary file S1: Text S6.

#### Text S7: Additional discussion of confounding factors

Some of the noise at lower taxonomic levels may putatively be attributable to the fact that we only constrained the topology at the phylum level, so lower taxonomic levels are certainly plagued by long-branch attraction artefacts [69,70].

A confounding factor in nematodes may be the fact that nematodes also exhibit an exceptionally complex evolutionary history of parasitism, with multiple origins of parasitic life histories [71], so we should account for the possibility that this complexity also confounds the signal.

We treated mitogenomes as a single marker due to the (mostly) absence of recombination, unilinear inheritance, and the fact that all PCGs are involved in the same (OXPHOS) pathway, but mitochondrial genes exhibit a relatively broad variability in evolutionary rates, commonly with *cox1* being the most conserved and *atp8* the least conserved [63]. To make sure that were are not capturing the signal from a minor proportion of the mitogenome, we analysed individual gene trees. We compared these trees to the one inferred using all 12 PCGs and selected the only comparatively large gene that exhibited a low correlation (*nad2*) to conduct pairwise analyses (Text S4 herein). As results were congruent with the result inferred using concatenated PCGs, this confirms that despite the notable intergenic variability in evolutionary rates, the effects of the two studied variables are consistent across individual genes.

While many categories used in the analyses exhibited a number of outliers (in terms of branch length), the distribution of outliers in higher-level datasets was such that their removal largely strengthened our conclusions (i.e. increased the probability of false negatives, and not false positives). We corroborated this by recreating our results using datasets with outliers removed.

As the demand for locomotion is also generally highly correlated with the mitochondrial abundance in locomotory muscles [72], it could be argued that species with reduced OXPHOS efficiency could simply compensate for this by increasing the number of mitochondria in their muscles. However, building and operating an increased number of mitochondria is a costly adaptation, so this would still affect the fitness of the individual. Therefore, mitogenomes with high energy production efficiency are more adaptive regardless of the number of mitochondria in muscle cells.

Episodic evolution may be an additional confounding factor. For example, the exceptionally long branch of parasitic flatworms is partly caused by the disproportionately long stem branch of Neodermata [21]. This implies that the transition to parasitism somewhere after the Cambrian explosion in this lineage was accompanied by a prolonged period of elevated evolutionary rates. This can affect a range of analyses.

#### Supplementary methods

##### Text S8: Datasets

The mitogenomes NC_059325, NC_059324, NC_054728, NC_053523, NC_050197, NC_046603, NC_044186, and NC_024698 were removed because the sequence was missing. We removed mitogenomes of several hybrids: NC_015838 (*Xenocypris davidi* x *Megalobrama amblycephala*), NC_028224 (*Megalobrama amblycephala* x *Megalobrama pellegrini*), NC_013995 (triploid

*Megalobrama amblycephala* x *Xenocypris davidi*), NC_013994 (diploid *Megalobrama amblycephala* x *Xenocypris davidi*), and NC_028224 (*Megalobrama amblycephala* x *Megalobrama pellegrini*). Several mitogenomes nominally belonging to different species were identical, which made us suspect species misidentification artefacts. In these cases we randomly removed one species. Identical pairs were: NC_031633 and NC_013074 (removed) (different genera), NC_047465 (removed) and NC_056102 (different genera), NC_040293 (removed) and NC_008534 (same genus), NC_024623 (removed) and NC_024645 (two hybrids with one conspecific parent), NC_020760 and NC_030175 (removed) (same genus), NC_036033 (removed) and NC_036034 (hybrids with same parent species), NC_015191 (removed) and NC_020011 (same genus), NC_026581 and NC_051547 (removed) (same genus), and NC_053682 and NC_053663 (removed) (same genus). We also removed several fragmented mitogenomes: *Anaticola crassicornis* NC_015998, *Liposcelis_entomophila* NC_025504 and NC_025503, and *Brachionus_plicatilis* NC_010472 and NC_010484. Finally, we removed NC_012980 because a conspecific (*Carassius auratus*) mitogenome already exists in the dataset.

We relied on the default GenBank taxonomic identity, retrieved from the NCBI's taxonomy database. As this is not regularly updated, there may be some minor differences with regard to more recent taxonomic changes. For example, in the NCBI's database, Acanthocephala and –Rotifera are two stand-alone phyla, but in recent classifications, they form a phylum Syndermata [73,74]. The dataset comprised all valid phyla aside from two small ones: Loricifera (37 species in total) and Micrognathozoa (only one known species).

As outgroups, we selected four most closely related phyla to Bilateria [75]: Ctenophora, Porifera, Cnidaria, Placozoa (two mitogenomes each). The eight species were: *Metridium senile* (NC_000933), *Acropora tenuis* (NC_003522), *Hoilungia* sp. (MT957399), *Polyplacotoma mediterranea* (NC_041549), *Axinella corrugata* (NC_006894), *Geodia_neptuni* (NC_006990), *Mnemiopsis leidyi* (NC_016117), *Coeloplana loyai* (LN898113).

#### Text S9: Classification of life-history (LH) strategies

Due to the high overall similarity between parasitoid and parasitic castrator strategies [76], the latter were also classified as parasitoids in most cases. Exceptions were some cestodes, which exhibit a mix of endoparasitic and parasitic castrator strategies [77]. With respect to our hypothesis (locomotory capacity, metabolic dependence on the host, and physical confinement to a single host) they are more similar to endoparasites than to parasitoids, so they were classified as endoparasites.

We assigned Oestridae to parasitoids, although their classification was somewhat ambiguous with respect to our hypotheses: larvae are endoparasitic, and adults are free-living, but adults generally do not feed during their short life. Therefore, the lineage could be treated as strictly parasitic as far as diet is concerned, but not as far as locomotory capacity and confinement to a single host are concerned.

A micropredator attacks more than one host (but one at a time), reducing each host's fitness by at least a small amount. Most micropredators are haematophagous (feeding on blood). They include annelids such as leeches, many insects (such as mosquitoes, tsetse flies and bed bugs), and even some vertebrates, such as lampreys and vampire bats. They often exhibit a mix of feeding strategies. For example, in many mosquitoes (Culicidae), typically both males and females feed non-parasitically. Only females need a blood meal in order to obtain the nutrients needed to produce eggs. Oxpecker *Buphagus erythrorhynchus* (Aves) is also such an example, their relationship with mammals was originally thought to be an example of mutualism, but more recent evidence suggests that oxpeckers may be facultative parasites (micropredators) [78].

Following the above definition, sheep ked *Melophagus ovinus* was classified as an ectoparasite and not as a micropredator, because it is believed to spend its entire life in the wool of a single sheep [79].

*Tinaminyssus melloi* and *Ptilonyssus chloris* (Arachnida: Mesostigmata: Rhinonyssidae) were classified as an endoparasites. Nasal mites show characteristics typical of endoparasitic species: reduced shielding, reduced setation, and overall, a body type that would have reduced mobility compared to free-living or ectoparasitic relatives [80].

*Trouessartia rubecula*, a feather mite, lives on birds' feathers, but feeds on feather oils, so we did not classify it as parasitic.

The glochidium (plural glochidia) is a microscopic larval stage of some freshwater mussels, aquatic bivalve molluscs in the families Unionidae and Margaritiferidae, the river mussels and European freshwater pearl mussels. This larva form has hooks, which enable it to attach itself to fish (for example to the gills of a fish host species) for a period before it detaches and falls to the substrate and takes on the typical form of a juvenile mussel. Since a fish is active and free-swimming, this process helps distribute the mussel species to areas that it could not reach otherwise. Despite its temporary attachment to the host, we did not classify any of these species as parasitic.

We failed to find sufficiently precise life-history information for *Goniophyto honshuensis, Miltogramma oestracea* (Sarcophagidae) and *Canthesancus helluo* (Reduviidae). As larvae of some Sarcophagidae are internal parasites of other insects such as Orthoptera, and some Reduviidae are blood-sucking ectoparasites, we did not assign an LH category to these three species, and we removed them from the dataset.

#### Text S10: Datasets used for selection tests

As the phylogenetic distribution of endoparasitic lineages employing code table 5 spanned three phyla, we tested a dataset comprising all endoparasitic and a selection of free-living species across nine phyla (N = 220). Ectoparasitic lineages in the table 5 dataset were limited to Arthropoda, so we selected a subdataset in the same way, with free-living lineages sampled only from Arthropoda (N = 178). There were too few endoparasitic species in the Arthropoda dataset (6), and they were included in the table 5 dataset, so we did not test them separately. However, we did test the Nematoda dataset (table 5) separately, as this phylum comprised a mix of endoparasitic and free-living species (the five EndoP/F species were excluded; N = 174). The only parasitic lineages employing other genetic codes (table 9) were limited to the phylum Platyhelminthes. Unfortunately, the number of free-living species in this dataset was rather lower (10), limiting the statistical power of the analyses. In our previous study, we used a larger Platyhelminthes dataset, comprising all available mitogenomes (not just the RefSeq dataset), which contained 30 free-living species.

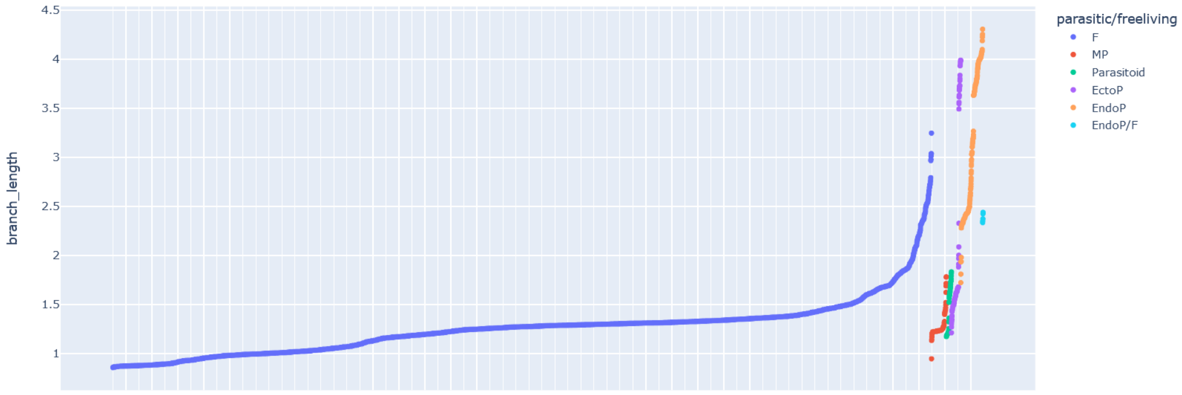

**Figure S12. Distribution of branch lengths across different life history categories.** Note: The branch length dataset had a nonnormal distribution (skewness = 3.49, kurtosis = 19.63, p<2.2E-16), but this is generally not a problem for large samples (>3000) [81]. Regardless, to reduce the magnitude of nonnormality we log-transformed the data. The distribution remained nonnormal, but with reduced values (skewness = 1.59, kurtosis = 7.83, p<2.2E-16).

#### References

1. Zou, H. *et al.* Inverted base composition skews and discontinuous mitochondrial genome architecture evolution in the Enoplea (Nematoda). *BMC Genomics* **23**, 376 (2022).
2. Zou, H. *et al.* Evolutionary rates of mitochondrial sequences and gene orders in Spirurina (Nematoda) are episodic but synchronised. *Water Biology and Security* **1**, 100033 (2022).
3. Zou, H. *et al.* The complete mitochondrial genome of parasitic nematode Camallanus cotti: extreme discontinuity in the rate of mitogenomic architecture evolution within the Chromadorea class. *BMC Genomics* **18**, 840 (2017).
4. Chen, F. *et al.* Sequencing of the Complete Mitochondrial Genome of Pingus sinensis (Spirurina: Quimperiidae): Gene Arrangements and Phylogenetic Implications. *Genes* **12**, 1772 (2021).
5. Zou, H. *et al.* The complete mitochondrial genome of Cymothoa indica has a highly rearranged gene order and clusters at the very base of the Isopoda clade. *PLOS ONE* **13**, e0203089 (2018).
6. Zou, H. *et al.* Architectural instability, inverted skews and mitochondrial phylogenomics of Isopoda: outgroup choice affects the long-branch attraction artefacts. *Royal Society Open Science* **7**, 191887 (2020).
7. Hua, C. J. *et al.* Basal position of two new complete mitochondrial genomes of parasitic Cymothoida (Crustacea: Isopoda) challenges the monophyly of the suborder and phylogeny of the entire order. *Parasites & Vectors* **11**, 628 (2018).
8. Zhang, D. *et al.* Mitochondrial genomes and 28S rDNA contradict the proposed obsoletion of the order Tetraonchidea (Platyhelminthes: Monogenea). *International Journal of Biological Macromolecules* **143**, 891–901 (2020).
9. Zhang, D. *et al.* Evidence for Adaptive Selection in the Mitogenome of a Mesoparasitic Monogenean Flatworm Enterogyrus malmbergi. *Genes* **10**, 863 (2019).

10. Zhang, D. *et al.* Mitochondrial genomes of two diplectanids (Platyhelminthes: Monogenea) expose paraphyly of the order Dactylogyridea and extensive tRNA gene rearrangements. *Parasites & Vectors* **11**, 601 (2018).

11. Zhang, D. *et al.* Three new Diplozoidae mitogenomes expose unusual compositional biases within the Monogenea class: implications for phylogenetic studies. *BMC Evolutionary Biology* **18**, 133 (2018).

12. Zhang, D. *et al.* Sequencing of the complete mitochondrial genome of a fish-parasitic flatworm Paratetraonchoides inermis (Platyhelminthes: Monogenea): tRNA gene arrangement reshuffling and implications for phylogeny. *Parasites & Vectors* **10**, 462 (2017).

13. Zhang, D. *et al.* Sequencing, characterization and phylogenomics of the complete mitochondrial genome of Dactylogyrus lamellatus (Monogenea: Dactylogyridae). *Journal of Helminthology* 1–12 (2017) doi:10.1017/S0022149X17000578.

14. Zhang, D. *et al.* Mitochondrial Genomes of Two Thaparocleidus Species (Platyhelminthes: Monogenea) Reveal the First rRNA Gene Rearrangement among the Neodermata. *International Journal of Molecular Sciences* **20**, 4214 (2019).

15. Zhang, D. *et al.* Homoplasy or plesiomorphy? Reconstruction of the evolutionary history of mitochondrial gene order rearrangements in the subphylum Neodermata. *International Journal for Parasitology* **49**, 819–829 (2019).

16. Li, W. X. *et al.* Characterization and phylogenomics of the complete mitochondrial genome of the polyzoic cestode Gangesia oligonchis (Platyhelminthes: Onchoproteocephalidea). *Journal of Helminthology* **94**, e58 (2020).

17. Li, W. X. *et al.* The complete mitochondrial DNA of three monozoic tapeworms in the Caryophyllidea: a mitogenomic perspective on the phylogeny of eucestodes. *Parasites & Vectors* **10**, 314 (2017).

18. Li, W. X. *et al.* Comparative mitogenomics supports synonymy of the genera Ligula and Digramma (Cestoda: Diphyllobothriidae). *Parasites & Vectors* **11**, 324 (2018).

19. Zou, H. *et al.* The complete mitochondrial genome of Gyrodactylus gurleyi (Platyhelminthes: Monogenea). *Mitochondrial DNA Part B* **1**, 383–385 (2016).
20. Zhang, D. *et al.* The complete mitochondrial genome of Gyrodactylus kobayashii (Platyhelminthes: Monogenea). *Mitochondrial DNA Part B* **1**, 146–147 (2016).
21. Jakovlić, I. *et al.* Drivers of interlineage variability in mitogenomic evolutionary rates in flatworms (Platyhelminthes) are multifactorial. 2022.09.11.507443 Preprint at https://doi.org/10.1101/2022.09.11.507443 (2022).
22. Jakovlić, I. *et al.* Slow crabs - fast genomes: locomotory capacity predicts skew magnitude in crustacean mitogenomes. *Molecular Ecology* **30**, 5488–5502 (2021).
23. Martin, A. P. & Palumbi, S. R. Body size, metabolic rate, generation time, and the molecular clock. *PNAS* **90**, 4087–4091 (1993).
24. Gillooly, J. F., Allen, A. P., West, G. B. & Brown, J. H. The rate of DNA evolution: Effects of body size and temperature on the molecular clock. *PNAS* **102**, 140–145 (2005).
25. Nabholz, B., Lanfear, R. & Fuchs, J. Body mass-corrected molecular rate for bird mitochondrial DNA. *Molecular Ecology* **25**, 4438–4449 (2016).
26. Thomas, J. A., Welch, J. J., Woolfit, M. & Bromham, L. There is no universal molecular clock for invertebrates, but rate variation does not scale with body size. *PNAS* **103**, 7366–7371 (2006).
27. Lanfear, R., Thomas, J. A., Welch, J. J., Brey, T. & Bromham, L. Metabolic rate does not calibrate the molecular clock. *PNAS* **104**, 15388–15393 (2007).
28. Thomas, J. A., Welch, J. J., Lanfear, R. & Bromham, L. A Generation Time Effect on the Rate of Molecular Evolution in Invertebrates. *Mol Biol Evol* **27**, 1173–1180 (2010).
29. Nabholz, B., Glémin, S. & Galtier, N. Strong Variations of Mitochondrial Mutation Rate across Mammals—the Longevity Hypothesis. *Mol Biol Evol* **25**, 120–130 (2008).
30. Galtier, N., Jobson, R. W., Nabholz, B., Glémin, S. & Blier, P. U. Mitochondrial whims: metabolic rate, longevity and the rate of molecular evolution. *Biology Letters* **5**, 413–416 (2009).

31. Welch, J. J., Bininda-Emonds, O. R. & Bromham, L. Correlates of substitution rate variation in mammalian protein-coding sequences. *BMC Evolutionary Biology* **8**, 53 (2008).
32. Moosmann, B. & Behl, C. Mitochondrially encoded cysteine predicts animal lifespan. *Aging Cell* **7**, 32–46 (2008).
33. Huang, D., Meier, R., Todd, P. A. & Chou, L. M. Slow Mitochondrial COI Sequence Evolution at the Base of the Metazoan Tree and Its Implications for DNA Barcoding. *J Mol Evol* **66**, 167–174 (2008).
34. Rand, D. M. Thermal habit, metabolic rate and the evolution of mitochondrial DNA. *Trends in Ecology & Evolution* **9**, 125–131 (1994).
35. Wang, Y. & Hekimi, S. Mitochondrial dysfunction and longevity in animals: Untangling the knot. *Science* **350**, 1204–1207 (2015).
36. Allio, R., Donega, S., Galtier, N. & Nabholz, B. Large variation in the ratio of mitochondrial to nuclear mutation rate across animals: implications for genetic diversity and the use of mitochondrial DNA as a molecular marker. *Mol Biol Evol* **34**, 2762–2772 (2017).
37. Saclier, N. *et al.* Life history traits impact the nuclear rate of substitution but not the mitochondrial rate in isopods. *Molecular Biology and Evolution* **35**, 2900–2912 (2018).
38. Galvani, A. P., Coleman, R. M. & Ferguson, N. M. The maintenance of sex in parasites. *Proceedings of the Royal Society of London. Series B: Biological Sciences* **270**, 19–28 (2003).
39. Jaenike, J. Mycophagous Drosophila and Their Nematode Parasites. *The American Naturalist* (1992) doi:10.1086/285365.
40. Lynch, M. The Evolution of Cladoceran Life Histories. *The Quarterly Review of Biology* **55**, 23–42 (1980).
41. Lynch, M., Koskella, B. & Schaack, S. Mutation pressure and the evolution of organelle genomic architecture. *Science* **311**, 1727–1730 (2006).
42. Lanfear, R., Ho, S. Y. W., Love, D. & Bromham, L. Mutation rate is linked to diversification in birds. *PNAS* **107**, 20423–20428 (2010).

43. Hardouin, E. a & Tautz, D. Increased mitochondrial mutation frequency after an island colonization: positive selection or accumulation of slightly deleterious mutations? *Biology Letters* **9**, 20121123–20121123 (2013).

44. James, J. E., Piganeau, G. & Eyre-Walker, A. The rate of adaptive evolution in animal mitochondria. *Mol Ecol* **25**, 67–78 (2016).

45. Bazin, E., Glémin, S. & Galtier, N. Population Size Does Not Influence Mitochondrial Genetic Diversity in Animals. *Science* **312**, 570–572 (2006).

46. Lynch, M. & Conery, J. S. The Origins of Genome Complexity. *Science* **302**, 1401–1404 (2003).

47. Palstra, F. P. & Ruzzante, D. E. Genetic estimates of contemporary effective population size: what can they tell us about the importance of genetic stochasticity for wild population persistence? *Molecular Ecology* **17**, 3428–3447 (2008).

48. Nabholz, B., Glémin, S. & Galtier, N. The erratic mitochondrial clock: variations of mutation rate, not population size, affect mtDNA diversity across birds and mammals. *BMC Evol Biol* **9**, 54 (2009).

49. Melvin, R. G. & Ballard, J. W. O. Cellular and population level processes influence the rate, accumulation and observed frequency of inherited and somatic mtDNA mutations. *Mutagenesis* **32**, 323–334 (2017).

50. Lajbner, Z., Pnini, R., Camus, M. F., Miller, J. & Dowling, D. K. Experimental evidence that thermal selection shapes mitochondrial genome evolution. *Scientific Reports* **8**, 9500 (2018).

51. Thomas, W. K. & Beckenbach, A. T. Variation in salmonid mitochondrial DNA: Evolutioinary constraints and mechanisms of substitution. *J Mol Evol* **29**, 233–245 (1989).

52. Lagisz, M., Poulin, R. & Nakagawa, S. You are where you live: parasitic nematode mitochondrial genome size is associated with the thermal environment generated by hosts. *Journal of Evolutionary Biology* **26**, 683–690 (2013).

53. Martin, A. P. Metabolic rate and directional nucleotide substitution in animal mitochondrial DNA. *Molecular Biology and Evolution* **12**, 1124–1131 (1995).

54. Faith, J. J. & Pollock, D. D. Likelihood Analysis of Asymmetrical Mutation Bias Gradients in Vertebrate Mitochondrial Genomes. *Genetics* **165**, 735–745 (2003).

55. Touchon, M., Arneodo, A., d'Aubenton-Carafa, Y. & Thermes, C. Transcription-coupled and splicing-coupled strand asymmetries in eukaryotic genomes. *Nucleic Acids Res* **32**, 4969–4978 (2004).

56. Britten, R. J. Rates of DNA sequence evolution differ between taxonomic groups. *Science* **231**, 1393–1398 (1986).

57. Christensen, A. C. Genes and Junk in Plant Mitochondria—Repair Mechanisms and Selection. *Genome Biol Evol* **6**, 1448–1453 (2014).

58. Lewis, S. C. *et al.* A Rolling Circle Replication Mechanism Produces Multimeric Lariats of Mitochondrial DNA in Caenorhabditis elegans. *PLOS Genetics* **11**, e1004985 (2015).

59. Oliveira, M. T., Haukka, J. & Kaguni, L. S. Evolution of the Metazoan Mitochondrial Replicase. *Genome Biol Evol* **7**, 943–959 (2015).

60. Weinstein, S. B. & Kuris, A. M. Independent origins of parasitism in Animalia. *Biology Letters* **12**, 20160324 (2016).

61. Dowton, M. & Austin, A. D. Increased genetic diversity in mitochondrial genes is correlated with the evolution of parasitism in the Hymenoptera. *J Mol Evol* **41**, 958–965 (1995).

62. Huyse, T., Poulin, R. & Théron, A. Speciation in parasites: a population genetics approach. *Trends in Parasitology* **21**, 469–475 (2005).

63. da Fonseca, R. R., Johnson, W. E., O'Brien, S. J., Ramos, M. J. & Antunes, A. The adaptive evolution of the mammalian mitochondrial genome. *BMC Genomics* **9**, 119 (2008).

64. Oliveira, D. C. S. G., Raychoudhury, R., Lavrov, D. V. & Werren, J. H. Rapidly Evolving Mitochondrial Genome and Directional Selection in Mitochondrial Genes in the Parasitic Wasp Nasonia (Hymenoptera: Pteromalidae). *Mol Biol Evol* **25**, 2167–2180 (2008).

65. Meiklejohn, C. D., Montooth, K. L. & Rand, D. M. Positive and negative selection on the mitochondrial genome. *Trends in Genetics* **23**, 259–263 (2007).

66. Keeling, P. J. *et al.* The Reduced Genome of the Parasitic Microsporidian Enterocytozoon bieneusi Lacks Genes for Core Carbon Metabolism. *Genome Biol Evol* **2**, 304–309 (2010).
67. Poulin, R. & Randhawa, H. S. Evolution of parasitism along convergent lines: from ecology to genomics. *Parasitology* **142**, S6–S15 (2015).
68. Slyusarev, G. S., Starunov, V. V., Bondarenko, A. S., Zorina, N. A. & Bondarenko, N. I. Extreme Genome and Nervous System Streamlining in the Invertebrate Parasite Intoshia variabili. *Current Biology* **30**, 1292-1298.e3 (2020).
69. Rota-Stabelli, O. *et al.* Ecdysozoan Mitogenomics: Evidence for a Common Origin of the Legged Invertebrates, the Panarthropoda. *Genome Biology and Evolution* **2**, 425–440 (2010).
70. Jakovlić, I. *et al.* Evolutionary History of Inversions in Directional Mutational Pressures in Crustacean Mitochondrial Genomes: Implications for Evolutionary Studies. *Molecular Phylogenetics and Evolution* **164**, 107288 (2021).
71. Blaxter, M. & Koutsovoulos, G. The evolution of parasitism in Nematoda. *Parasitology* **142**, S26–S39 (2015).
72. Childress, J. J. Are there physiological and biochemical adaptations of metabolism in deep-sea animals? *Trends in Ecology & Evolution* **10**, 30–36 (1995).
73. Sielaff, M. *et al.* Phylogeny of Syndermata (syn. Rotifera): Mitochondrial gene order verifies epizoic Seisonidea as sister to endoparasitic Acanthocephala within monophyletic Hemirotifera. *Molecular Phylogenetics and Evolution* **96**, 79–92 (2016).
74. Monks, S. Phylogeny of the Acanthocephala based on morphological characters. *Syst Parasitol* **48**, 81–115 (2001).
75. Laumer, C. E. *et al.* Revisiting metazoan phylogeny with genomic sampling of all phyla. *Proceedings of the Royal Society B: Biological Sciences* **286**, 20190831 (2019).
76. Kuris, A. M. Trophic Interactions: Similarity of Parasitic Castrators to Parasitoids. *The Quarterly Review of Biology* **49**, 129–148 (1974).

77. Arnott, S. A., Barber, I. & Huntingford, F. A. Parasite–associated growth enhancement in a fish–cestode system. *Proceedings of the Royal Society of London. Series B: Biological Sciences* **267**, 657–663 (2000).
78. Plantan, T., Howitt, M., Kotzé, A. & Gaines, M. Feeding preferences of the red-billed oxpecker, Buphagus erythrorhynchus: a parasitic mutualist? *African Journal of Ecology* **51**, 325–336 (2013).
79. Small, R. W. A review of Melophagus ovinus (L.), the sheep ked. *Veterinary Parasitology* **130**, 141–155 (2005).
80. Hilario-Pérez, A. D. & Dowling, A. P. G. Nasal mites from specimens of the brown-headed cowbird (Icteridae: Molothrus ater) from Texas and Arkansas, U.S.A. *Acarologia* **58**, 296–301 (2018).
81. Li, X., Wong, W., Lamoureux, E. L. & Wong, T. Y. Are Linear Regression Techniques Appropriate for Analysis When the Dependent (Outcome) Variable Is Not Normally Distributed? *Investigative Ophthalmology & Visual Science* **53**, 3082–3083 (2012).
